# A molecular and spinal circuit basis for the functional segregation of itch and pain

**DOI:** 10.1101/2025.07.31.667966

**Authors:** Myung-chul Noh, Kelly A. Corrigan, Sean-Paul G. Williams, Cedric Peirs, Michael J. Leone, Daniel J. Headrick, Mustafa Guvercin, Suhjin Lee, BaDoi N. Phan, Deepika Yeramosu, Sudhagar Babu, Ashley R. Brown, Robert Van De Weerd, Xinyi Zhao, Richard P. Dum, Hansruedi Mathys, Andreas R. Pfenning, Rebecca P. Seal

**Affiliations:** Department of Neurobiology, University of Pittsburgh; Pittsburgh, 15260, United States; Pittsburgh Center for Pain Research, University of Pittsburgh; Pittsburgh, 15260, United States; Medical Scientist Training Program, University of Pittsburgh; Pittsburgh, 15260, United States; Computational Biology Department, School of Computer Science, Carnegie Mellon University; Pittsburgh, 15213, United States; Univ. Clermont-Auvergne, INSERM U1107: Neuro-Dol; Clermont Ferrand, F-63001, France

## Abstract

Recent advances reveal an extensive cellular diversity within the dorsal horn. How this complexity processes distinct sensations, like itch and pain, remains a fundamental question. We discovered hidden within a population of neurons expressing the gastrin-releasing peptide receptor (*Grpr*+), thought to be itch-specific, are highly homologous yet functionally distinct subtypes distinguished by expression of Tachykinin-1 (*Tac1*). While the *Tac1*- subtype mediates itch, the *Tac1*+ subtype mediates mechanical allodynia across diverse pain states. Inhibitory populations and differential sensitivities to GRP serve as key modulators of the *Grpr*+ neuron subtypes, shaping modality specific output. Leveraging computationally designed genomic enhancers to silence *the Tac1*- population reverses itch while silencing the *Tac1+* subtype reverses mechanical allodynia broadly. The work demonstrates the nuance of differential sensory modality coding within the dorsal horn and the power of genomic enhancer-based strategies for modality-specific targeting.

## Main Text

Chronic pain represents a major unmet clinical need. Mechanical allodynia is a common and debilitating form; whereby innocuous touch evokes severe pain (*1*). Arising from maladaptive plasticity within the dorsal horn, diverse aetiologies like nerve trauma, diabetes, postherpetic neuralgia, and multiple sclerosis can cause mechanical allodynia (*1–4*). The dorsal horn also receives and processes an array of stimuli, innocuous and noxious temperatures, diverse mechanical inputs, and distinct forms of itch, raising fundamental questions about how its circuit organization maintains the fidelity of qualitatively distinct sensations, including chronic pain (*2–5*). The enormous challenge of deciphering the organization is further underscored by a prevalence of nociceptors (*6–12*) spinal interneurons (*13–16*) and projection neurons (*17–19*) showing polymodal response properties, though supporting evidence is largely correlative rather than causal. Nevertheless, understanding the functional organization of the dorsal horn and how it adapts or becomes maladaptive during chronic pain states, including with respect to different aetiologies, are key goals for the field and for developing safe and efficacious therapeutic interventions.

Gaps in knowledge regarding the functional organization of the dorsal horn stem in part from the highly heterogeneous nature of the neurons. Although well characterized in terms of their spatial distributions, morphological and electrophysiological properties (*20, 21*), and neurochemistry (*22, 23*), a coherent functional organization has not emerged from these datasets. Recent transcriptomic studies have provided an increasingly refined, species-integrated classification of dorsal horn neuron subtypes from mice to humans, thus enabling more precise functional analyses and enhancing translational potential (*24–27*). However, linking these transcriptomically defined subtypes to specific functions in somatosensory processing, particularly in chronic pain states, remains a pressing challenge. An immediate advancement that makes this feasible is the development of cell-type specific enhancers (*28–30*), that enable the required genetic access and precision to dissect the complex circuits of the dorsal horn.

Based on observations of both single and polymodality coding of somatosensory information within the neuroaxis, two major models have been described: one posits dedicated circuits for each sensation, while the other proposes differential activity patterns across shared neuronal populations (*5*). The models apply not only to acute somatosensory modalities but also persistent pain states. While initial models of mechanical allodynia proposed a single dorsally-directed pathway from lamina III to the lamina I projection neurons, recent studies reveal a network of overlapping circuits differentially engaged depending on the nature of the aetiology (*31, 32*). Despite the significant progress made in identifying key neurons and pathways specific to mechanical hypersensitivity induced by a wide range of inflammatory and neuropathic injuries (*31, 32*), the circuits important following polyneuropathic injuries, for example in diabetes, remain largely unknown.

### Distinct dorsal horn circuits mediate polyneuropathic pain

Polyneuropathy, characterized by widespread, symmetrical damage to multiple peripheral nerves, is a highly prevalent subtype of neuropathic injury (*33*). Common causes include diabetes, which affects hundreds of million people globally (*33*) and chemotherapy, where the resulting polyneuropathy affects a substantial portion of patients and often limits the use of these life-saving cancer treatments (*34*). Given that polyneuropathy shares aetiological features with other neuropathic injuries that cause damage to the peripheral nervous system, we initially hypothesized that the spinal circuits mediating mechanical allodynia would be similar. We therefore examined the role of neuronal populations known to be important for mechanical allodynia in neuropathic and inflammatory injury models. Strikingly, and contrary to our initial expectations, our experiments reveal that these known circuits are dispensable for mechanical allodynia in established models of polyneuropathic injury.

Specifically, we tested whether the vesicular glutamate transporter 3 (VGLUT3) has a role in the oxaliplatin model of chemotherapeutic polyneuropathic pain, long-course streptozotocin (STZ) chemical (type I) and leptin*^ob/ob^* genetic (type II) models of diabetic polyneuropathic pain using VGLUT3 knockout (KO) mice. Previously, it was demonstrated that the KO mice have significantly attenuated mechanical allodynia compared to wildtype (WT) littermates in a range of inflammatory and neuropathic injuries and that this phenotype is caused by the loss of VGLUT3 from a population of dorsal horn neurons during development (*31*). Surprisingly, the KO mice show normal mechanical allodynia in all three polyneuropathic pain models (fig. S1A). Next, we tested the role of the cholecystokinin (CCK), protein kinase C gamma (PKCγ), and calretinin (CR) neurons, which were shown to be important for mechanical allodynia in neuropathic and/ or inflammatory injuries using chemogenetic inhibition (*32*). Using this same approach, we show that these neuronal populations are dispensable for mechanical allodynia induced by the long course STZ model of diabetic polyneuropathic pain (fig. S1, C to E).

### *Grpr^+^* neurons mediate mechanical allodynia

Given these unexpected findings, we next turned our attention to a long-studied population of dorsal horn neurons called vertical cells, which had been implicated in mechanical allodynia but whose molecular identity was not known. Within the complex circuity of the dorsal horn, vertical cells represent a morphologically distinct neuronal population characterized by cell bodies in outer lamina II and dendrites extending ventrally into lamina III (*20, 35*). Vertical cells are uniquely positioned to integrate diverse inputs, including from nociceptive and low-threshold mechanoreceptor afferents, as well as local interneurons (*15, 36, 37*). This, coupled with their direct axonal contacts onto lamina I projection neurons (*38*), places vertical cells in an ideal position to modulate pain signalling. Consistent with a role for these cells in mechanical allodynia, it was demonstrated that peripheral injury-induced spinal disinhibition allows low-threshold mechanosensory inputs to engage the vertical cells (*37, 39*). Recent work identified *Grpr-*expressing neurons as having properties consistent with this type of vertical cell (*14*), including monosynaptic inputs from both excitatory and inhibitory spinal interneurons (fig. S2, F to G and I to J) (*40, 41*), a broad range of primary sensory afferents (fig. S2, H and K) (*14, 42, 43*) as well as axonal contacts directly with projection neurons in lamina I and the lateral spinal nucleus (*43, 44*). While *Grpr^+^*neurons exhibit anatomical features of vertical cells previously implicated in pain modulation, they were reported to be essential only for itch and to not have a role in mechanical allodynia induced by inflammatory and neuropathic injuries (*45, 46*). However, a more recent study showed that chemogenetic activation of these neurons not only elicited biting behaviour, consistent with itch, but also caused licking and guarding behaviour suggestive of pain (*14*). We therefore re-examined the role of *Grpr^+^* neurons in mechanical allodynia across a broad range of injury types.

As an initial test for a role of the *Grpr* neurons in mechanical allodynia, we used the long-course STZ model of diabetic polyneuropathic pain and measured the upregulation of *cFos*, a marker of neuronal activity, in the superficial dorsal horn (Fig. 1, A to C). In these experiments, mice were walked on a treadmill to provide a low threshold mechanical stimulus. As expected, the STZ-treated mice, which exhibited mechanical allodynia behaviour (fig. S1), showed significantly more *cFos^+^*cells in the superficial dorsal horn compared to naïve mice (Fig. 1D). Importantly, the percentage of *Grpr^+^* cells expressing *cFos* was increased significantly in STZ mice compared to naïve mice (Fig. 1E), and the percentage of *cFos^+^* cells expressing *Grpr* (Fig. 1F) was also significantly greater in the STZ mice. These results provide support for the hypothesis that *Grpr^+^* neurons are involved in the mechanical allodynia induced by polyneuropathic injury.

**Fig. 1.**
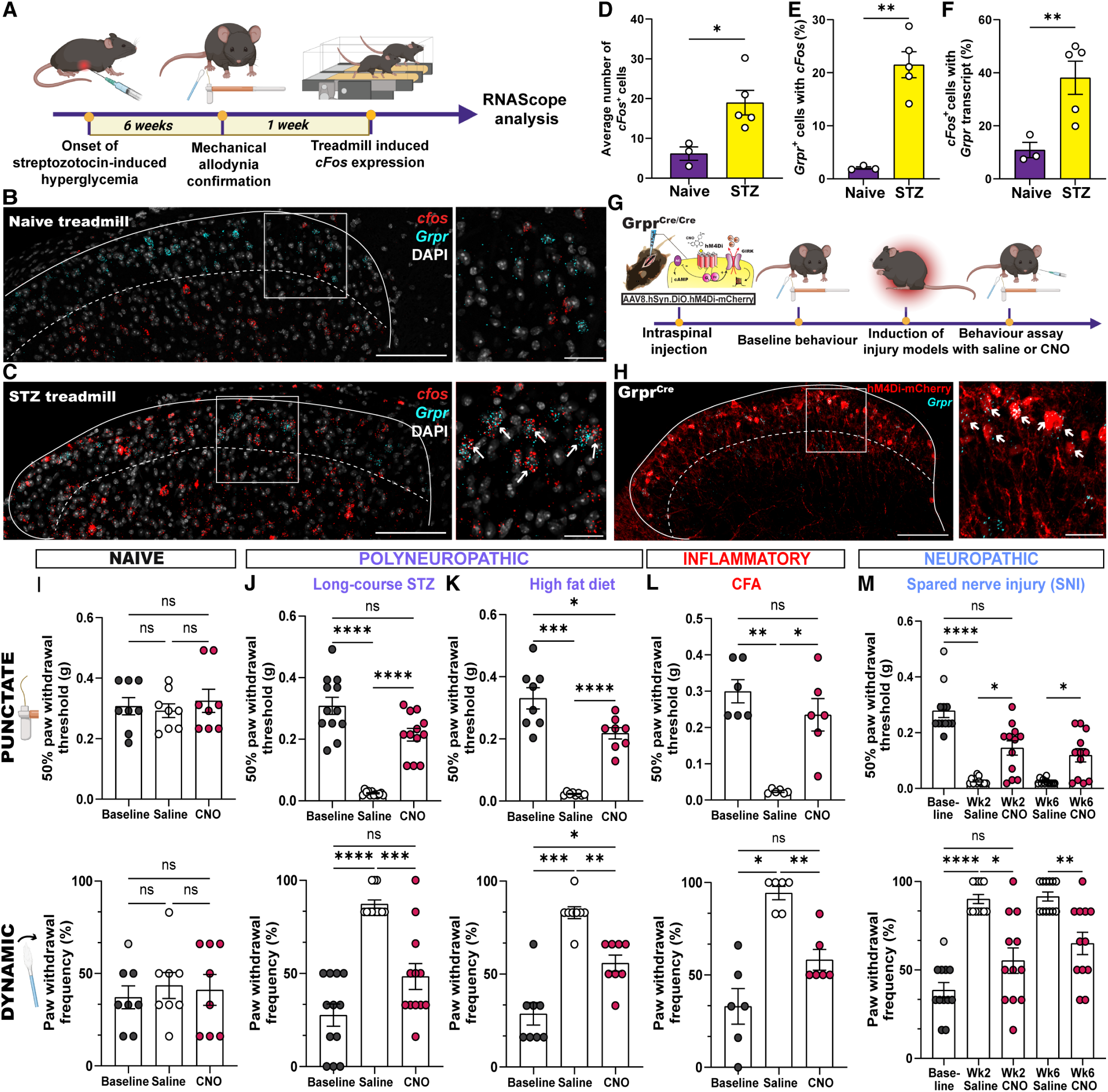
*Grpr^+^* neurons mediate mechanical allodynia across multiple injury types. **(A)** Experimental strategy to examine *cFos* expression in the dorsal horn. **(B and C)** Transverse lumbar spinal dorsal horn section showing *cFos* (red) and *Grpr* (cyan) mRNA expression in (B) naïve and (C) long-course streptozotocin (STZ) mice following treadmill stimulation. Arrows indicate cells co-expressing *cFos* and *Grpr*. (**D to F)** Quantification of *cFos^+^* cells in the superficial dorsal horn of naïve (n=3) and long-course STZ mice (n=5). (D) Number of *cFos^+^* cells. (E) Percentage of *Grpr^+^*cells with *cFos.* (F) Percentage of *cFos^+^* cells that are *Grpr^+^*. **(G)** Experimental strategy to examine the role of spinal *Grpr^+^* neurons in mechanical allodynia. **(H)** Lumbar dorsal horn section from a homozygous Grpr^Cre^ mouse showing Cre-dependent hm4di-mcherry expression (red) and *Grpr* transcript (cyan) after intraspinal delivery of AAV8.hSyn.DIO.hm4di-mCherry. Arrows indicate cells with hM4Di-mCherry co-expressing *Grpr* transcript (n=3 mice). **(I to M)** Mechanical sensitivity assessment in Grpr^Cre^ mice expressing hM4Di. Top panels depict 50% paw withdrawal thresholds for punctate mechanical sensitivity (von Frey test). Bottom panels depict paw withdrawal frequency for dynamic mechanical sensitivity (cotton swab test). (I) Uninjured mice (n=8). (J) Mice with diabetic polyneuropathy induced by long-course STZ model (n=12). (K) Mice with diabetic polyneuropathy induced by high fat diet (n=8). (L), Mice with inflammatory injury induced by complete Freund’s adjuvant (CFA) (n=6). (M) Mice with neuropathic injury induced by spared nerve injury (SNI) (n=12). Dashed lines in representative figures (B, C and H) delineate lamina II/III border. Scale bars, 100 μm (main images) and 30 μm (insets). Data are presented as mean ± s.e.m. Statistical analysis was performed with unpaired two-tailed Welch’s t-test (D to F), and repeated-measures one-way ANOVA with Bonferroni’s post-hoc test (I to M). *P < 0.05, **P < 0.01, ***P < 0.001, ****P < 0.0001.

To causally test whether *Grpr^+^* neurons are important for mechanical allodynia induced by polyneuropathic injuries, we used the chemogenetic approach to acutely silence *Grpr^+^* neurons while testing pain behaviour (Fig. 1G). The inhibitory DREADD (hM4Di) receptor (*47*) was targeted specifically to dorsal horn *Grpr^+^* neurons by injecting AAV8.hSyn.DIO.hM4Di-mCherry into the lumbar dorsal horn of homozygous Grpr^Cre^ mice (Fig 1H; 93.01 ± 1.043% of hM4Di^+^ cells were *Grpr^+^*; n=3 mice). Initially, we tested whether silencing *Grpr^+^* neurons with the hM4Di ligand, clozapine-N-oxide (CNO), affects baseline punctate (von Frey) or dynamic (cotton swab) mechanical thresholds. Grpr^hM4Di^ mice injected with CNO (5 mg kg^-1^, i.p) showed similar mechanical thresholds as saline injected mice (Fig. 1I). These results are in line with previous findings where baseline mechanosensory thresholds were not altered by ablating *Grpr^+^* neurons using intrathecally delivered bombesin-saporin (*46*). We next tested the impact of silencing the *Grpr^+^* neurons in the two diabetic polyneuropathy models, the type 1 diabetes model, *i.e.* long-course STZ, and the type II diabetes model, *i.e.* high fat diet (HFD) (fig. S3). In both injury models, CNO-mediated silencing significantly attenuated punctate and dynamic allodynia (Fig. 1, J and K). Despite previous reports, which relied on bombesin-saporin ablation (*46*) and showed that *Grpr^+^* neurons are dispensable for the mechanical allodynia induced by inflammatory and neuropathic injuries, we instead find that mechanical thresholds return to baseline when *Grpr^+^* neurons are silenced following induction of both the complete Freund’s adjuvant (CFA) inflammatory model and the spared nerve injury (SNI) neuropathic model (Fig. 1, L and M). Together, the results provide strong evidence that *Grpr^+^* neurons are in fact important for mechanical allodynia induced by all three types of injuries.

### *Grpr^+^* neurons are transcriptomically heterogeneous

These behavioural results indicate a convergence of itch and pain processing within a single neuronal population defined by the expression of *Grpr*, which raises a fundamental question about how the circuitry for these different modalities is organized within the spinal dorsal horn. We reasoned that *Grpr^+^*neurons participate in the circuitry for itch and mechanical allodynia in one of two ways: 1) both modalities are differentially encoded by the same *Grpr*^+^ neurons; or 2) the *Grpr^+^* neurons are not a homogenous population but rather are composed of functionally and transcriptomically distinct subpopulations that separately encode itch and pain. To investigate the latter possibility, we examined the single-nucleus RNA-sequencing (snRNA-seq) dataset of a meta-analysis performed on six mouse spinal cord studies (*25*). In this dataset, *Grpr^+^* nuclei were distributed across three excitatory neuron clusters, Excit-14, 15, and 16 (fig. S4a). Interestingly, the Excit-14 cluster showed high levels of *Grpr* transcript while the Excit-16 cluster trended towards lower levels of *Grpr* transcript (fig. S4, B and C). Moreover, we identified a marker gene, tachykinin 1 (*Tac1*), that clearly differentiates these two major subpopulations, allowing us to examine their individual contributions to itch and pain processing. Specifically, the differential gene expression analysis revealed a striking enrichment of *Tac1* expression within *Grpr^+^* nuclei of Excit-16 (90% of the *Grpr^+^* nuclei) compared to the *Grpr^+^*nuclei of Excit-14 (14% of the *Grpr^+^* nuclei) (fig. S4, B to D).

As a first step to identify potential functional differences between the two major *Grpr^+^* subpopulations (Excit-14 and Excit-16), we examined their spatial distribution in the lumbar dorsal horn using fluorescence *in situ* hybridization (FISH) (Fig. 2A). *Grpr*^+^ neurons that lack *Tac1* (*Grpr*^+^*Tac1*^-^) are predominantly located in the lamina I and outer layer of lamina II, while those that co-express *Tac1* (*Grpr*^+^*Tac1*^+^) reside in both the outer and inner layer of lamina II (Fig. 2A). Using spatial transcriptomics analysis, we further examined the distribution of the two major *Grpr*^+^ subpopulations using our recently generated species-integrated atlas of dorsal horn neuron subtypes (Fig. 2B and fig.S5) (*26*). The cross-species neuron subtypes defined in this atlas were generated by integrating individual neuronal nuclei from the mouse meta-analysis with neuronal nuclei from macaque and human RNA-seq datasets (*26*). Here, we find the vast majority of *Grpr^+^* cells (∼85%) are divided between the excitatory neuron subtypes Exc-SKOR2 and Exc-LMO3 (Fig. 2, B and C). Indeed, mouse subtypes Excit-14 and Excit-16 correspond to the species conserved subtypes, Exc-SKOR2 and Exc-LMO3, respectively (*26*). Spatial transcriptomic analysis of mouse lumbar dorsal horn slices also reveals similar relationships between the neuron subtypes and the levels of *Grpr* and *Tac1* transcripts. Specifically, the number of *Grpr* transcripts per cell are highest in *Grpr^+^* neurons of the Exc-SKOR2 subtype and trend lower in neurons of the Exc-LMO3 subtype, while the number of *Tac1* transcripts per cell are higher in *Grpr^+^* neurons of the Exc-LMO3 subtype and are essentially negligible in neurons of the Exc-SKOR2 subtype (Fig. 2D). Further analysis of the cross-species neuron subtype distribution of *Grpr*^+^ neurons shows 73% *of Grpr^+^Tac1*^-^ neurons belong to the Exc-SKOR2 subtype while a large majority (∼82%) of the *Grpr*^+^ *Tac1*^+^ neurons belong to the Exc-LMO3 subtype (Fig. 2E). With respect to the distribution of the two major *Grpr*^+^ subpopulations as a percentage of total neurons within each subtype, *Grpr^+^Tac1*^-^ neurons comprise ∼63% of the neurons in Exc-SKOR2 while *Grpr^+^Tac1*^+^ neurons comprise ∼23% of the neurons in Exc-LMO3 (Fig. 2F).

**Fig. 2:**
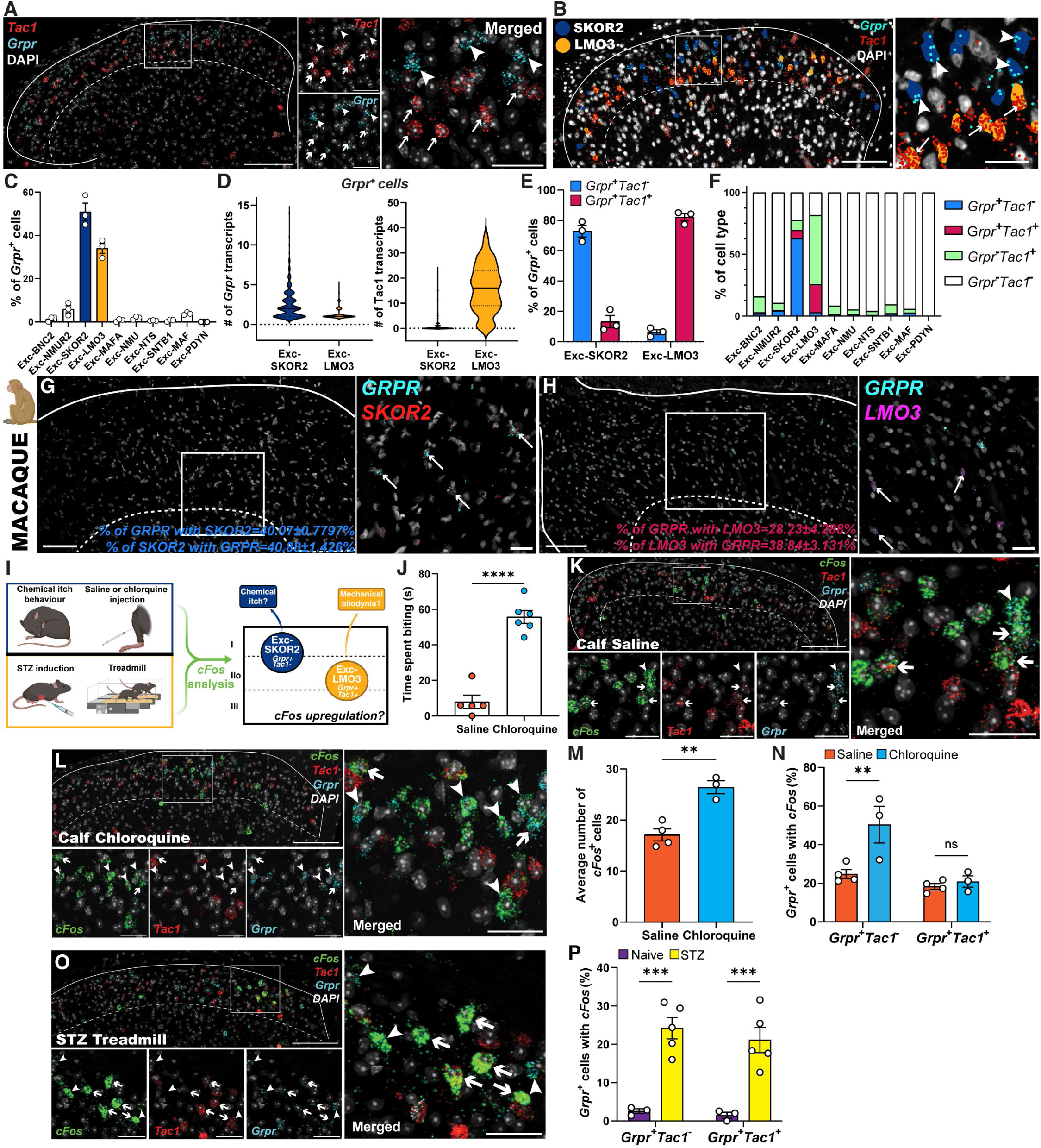
Differential activation of transcriptomically distinct *Grpr^+^* neuron subtypes to chemical itch and mechanical allodynia. **(A)** Lumbar dorsal horn section of a mouse showing *Tac1* (red) and *Grpr* (cyan) transcripts. Arrowheads indicate *Grpr^+^Tac1^-^* cells and arrows indicate *Grpr^+^Tac1^+^*cells. **(B)** Xenium spatial transcriptomics image of lumbar dorsal horn section of a mouse showing Exc-SKOR2 (blue) and Exc-LMO3 (yellow) neurons with *Grpr* (cyan) and *Tac1* (red) transcripts. Arrowheads indicate Exc-SKOR2 cells expressing only *Grpr*. Arrows indicate Exc-LMO3 cells expressing *Grpr* and *Tac1*. **(C)** Distribution of *Grpr^+^* cells across species-conserved dorsal horn excitatory neuron populations (n=3 mice). **(D)** Violin plots showing *Grpr* (left) and *Tac1* (right) transcript counts within *Grpr^+^*cells in Exc-SKOR2 and Exc-LMO3 populations (n=3 mice) Solid lines: median; dashed lines: quartiles. **(E)** Distribution of *Grpr^+^Tac1^-^* (blue) and *Grpr^+^Tac1^+^* (magenta) cells across Exc-SKOR2 and Exc-LMO3 populations (n=3 mice). **(F)** Relative proportion of *Grpr^+^Tac1^-^* (blue), *Grpr^+^Tac1^+^* (magenta), *Grpr^-^ Tac1^+^* (green), and *Grpr^-^ Tac1^-^* (white) neurons across species-conserved dorsal horn excitatory neuron populations (n=3 mice). **(G to H)** Lumbar dorsal horn section of a macaque showing *GRPR* (cyan), *SKOR2* (red), and *LMO3* (purple) transcripts. Arrows indicate *GRPR^+^* cells expressing (G) *SKOR2* or (H) *LMO3* (n=2 macaques). **(I)** Experimental strategy to examine *cFos* upregulation in Exc-SKOR2 or Exc-LMO3 neurons following mechanical stimulation (before and after polyneuropathic injury) and intradermal calf injection of saline or chloroquine. **(J)** Itch-like behaviour (time spent biting calf) following intradermal injection of saline (n=5) or chloroquine (n=6). **(K and L)** Lumbar dorsal horn section showing *cFos* (green), *Tac1* (red), and *Grpr* (cyan) transcripts after intradermal (K) saline or (L) chloroquine injections. Arrowheads indicate *cFos* expression in *Grpr^+^Tac1^-^* cells. Arrows indicate *cFos* expression in *Grpr^+^Tac1^+^* cells. **(M and N)** Quantification of *cFos^+^* cells in lumbar dorsal horn (n=4 mice saline, n=3 mice chloroquine). (M) Average number of *cFos^+^*cells. (N) Percentage of *Grpr^+^* cells expressing *cFos*. **(O)** Lumbar dorsal horn section showing *cFos* (green), *Tac1* (red), and *Grpr* (cyan) transcripts in a long-course STZ mouse after treadmill stimulation. Arrowheads indicate *cFos* expression in *Grpr^+^Tac1^-^* cells. Arrows indicate *cFos* expression in *Grpr^+^Tac1^+^* cells. **(P)** Percentage of *Grpr^+^* cells expressing *cFos* transcript in naïve (n=3 mice) and long-course STZ (n=5 mice) mice after treadmill stimulation (same mice as Fig. 1D to F). Data are mean ± s.e.m. unless stated otherwise. Dashed lines in representative figures (A, B, G, and K to O) delineate lamina II/III border. Scale bars: 100 μm (main images) and 30 μm (insets). Statistical analysis was performed with unpaired two-tailed Welch’s t-test (J and M), and two-way ANOVA with Bonferroni’s post-hoc test (N and P). *P < 0.05, **P < 0.01, ***P < 0.001, ****P < 0.0001

Lastly, we performed FISH in the lumbar dorsal horn of macaques to assess *GRPR* transcript expression in Exc-SKOR2 (*SKOR2*^+^) and Exc-LMO3 (*LMO3^+^*) subtypes (Fig. 2, G and H). We found that the expression of *GRPR* transcript was highly conserved within these subtypes. In the macaque dorsal horn, as in mice, ∼40% of *GRPR*^+^ cells co-expressed SKOR2, and about 40% of Exc-SKOR2 neurons expressed *GRPR* (Fig. 2G). Likewise, ∼28% of *GRPR^+^* cells co-expressed LMO3, while about 39% of Exc-LMO3 neurons expressed *GRPR* (Fig. 2H). These analyses therefore identify two major *Grpr^+^* neuron subpopulations: a *Grpr^+^Tac1^-^*population, characterized by relatively higher *Grpr* transcript levels, belonging primarily to the species-conserved Exc-SKOR2 subtype, and the *Grpr^+^Tac1^+^* population, characterized by relatively lower *Grpr* transcript levels but high *Tac1* transcript levels, corresponding primarily to the Exc-LMO3 subtype.

### *Grpr^+^* neuron subtypes respond differentially to itch and pain

The distinct transcriptomic profiles and spatial distributions of the two major *Grpr^+^* subpopulations suggest they may also differ in their functional roles and local circuit interactions. To assess the functional roles, we first examined whether the two subpopulations are differentially activated by pruritogens and mechanical allodynia by performing *cFos* analyses (Fig. 2I). Because chemical itch induced by chloroquine is known to be conveyed by neurons that reside within dorsal horn laminae I and II outer (*48, 49*), and the *Grpr^+^Tac1^-^* neurons are predominantly located within this region, we hypothesized that chloroquine will preferentially activate this subpopulation. As expected, itch-like behaviour and the total number of *cFos^+^* neurons in the superficial dorsal horn are increased significantly following injection of chloroquine compared to saline (Fig. 2, J to M). Importantly, *cFos* is significantly upregulated only in *Grpr^+^Tac1^-^* neurons following injection of chloroquine compared to saline injection (Fig. 2, K, L, and N). The latter treatment also controls for the needlestick, which causes *cFos* expression in both subpopulations evenly (Fig. 2, K, L, and N). With respect to mechanical allodynia, mice treated with long-course STZ and walked on a treadmill show an equally significant increase in *cFos* expression in both subpopulations of *Grpr^+^* neurons compared to uninjured mice (Fig. 2, O and P). This interestingly mirrors the pattern of *cFos* upregulation we observe with needlestick. These findings suggest that *Grpr^+^Tac1^-^* neurons preferentially mediate chemical itch, whereas the specific contributions of *Grpr^+^* neuron subtypes to mechanical allodynia remain unclear.

### Differential inhibitory modulation of *Grpr^+^* neuron subtypes

The lack of clarity regarding the roles of the two *Grpr^+^* neuron subpopulations in mechanical allodynia prompted us to investigate whether distinct inhibitory interneuron populations differentially regulate them. This line of investigation is based on previous reports indicating that parvalbumin (PV) interneurons in the dorsal horn impact mechanical allodynia but not chemical itch (*37, 41, 50*), while prodynorphin (PDYN) neurons can impact both itch (*41, 51, 52*) and pain (*15, 53*). Specifically, spinal inhibitory PV (iPV) interneurons have been implicated in gating mechanical allodynia, in part through direct inhibitory input onto vertical cells (of unknown molecular identity, which we now hypothesize are *Grpr^+^*neurons) and onto the Aβ afferent terminals that directly innervate these vertical cells (*37, 50, 54*). Conversely, a separate study showed broad chemogenetic activation of PV interneurons, which also includes excitatory neurons (*54*), had no effect on itch-like behaviours (*41*). On the other hand, PDYN interneurons have been implicated in modulating chemical itch through the direct inhibition of *Grpr^+^* neurons (*41, 51, 52*). PDYN expressing interneurons have also been implicated in modulating mechanical allodynia (*15, 53*). In one study, PDYN-lineage neurons were ablated with diphtheria toxin in the adult and in a second study, dynorphin lineage neurons were broadly inhibited. In both studies, the manipulations induced mechanical hypersensitivity (*15, 53*). Furthermore, in a third study, activation of PDYN neurons, which includes both excitatory and inhibitory neurons, produced mechanical hypersensitivity (*52*), indicating a need for a more specific manipulation of just the inhibitory PDYN neurons (iPDYN) to dissect the circuitry. Taking the previous studies into consideration, we examined whether the two inhibitory interneuron populations are directly presynaptic to *Grpr^+^* neurons using monosynaptic retrograde rabies tracing. Indeed, injection of Grpr^Cre^ mice with a Cre-dependent AAV helper virus, followed by the injection of EnvA G-deleted rabies encoding a dsRedXpress reporter, led to expression of the reporter in both iPV and iPDYN interneurons (∼13% and ∼6% of dsRedXpress, respectively; fig. S2, F, G, and J).

We then tested whether the two inhibitory interneuron populations differentially modulate itch-like behaviour and mechanical allodynia and combined these behavioural tests with the analysis of *cFos* expression in the two major *Grpr^+^* subpopulations (Fig. 3A). We hypothesized that if the activation of iPV neurons does not impact itch-like behaviour, it will also not supress *cFos* upregulation in the *Grpr^+^Tac1^-^* subpopulation. We also hypothesized that if activation of iPV neurons attenuates mechanical allodynia, then it will also significantly reduce the *cFos* normally upregulated by mechanical allodynia in the *Grpr^+^Tac1^+^* subpopulation or in both subpopulations. Conversely, we hypothesized that activation of iPDYN neurons will reduce both itch-like behaviour and mechanical allodynia leading to attenuation of *cFos* expression in both *Grpr^+^* subpopulations.

**Fig. 3:**
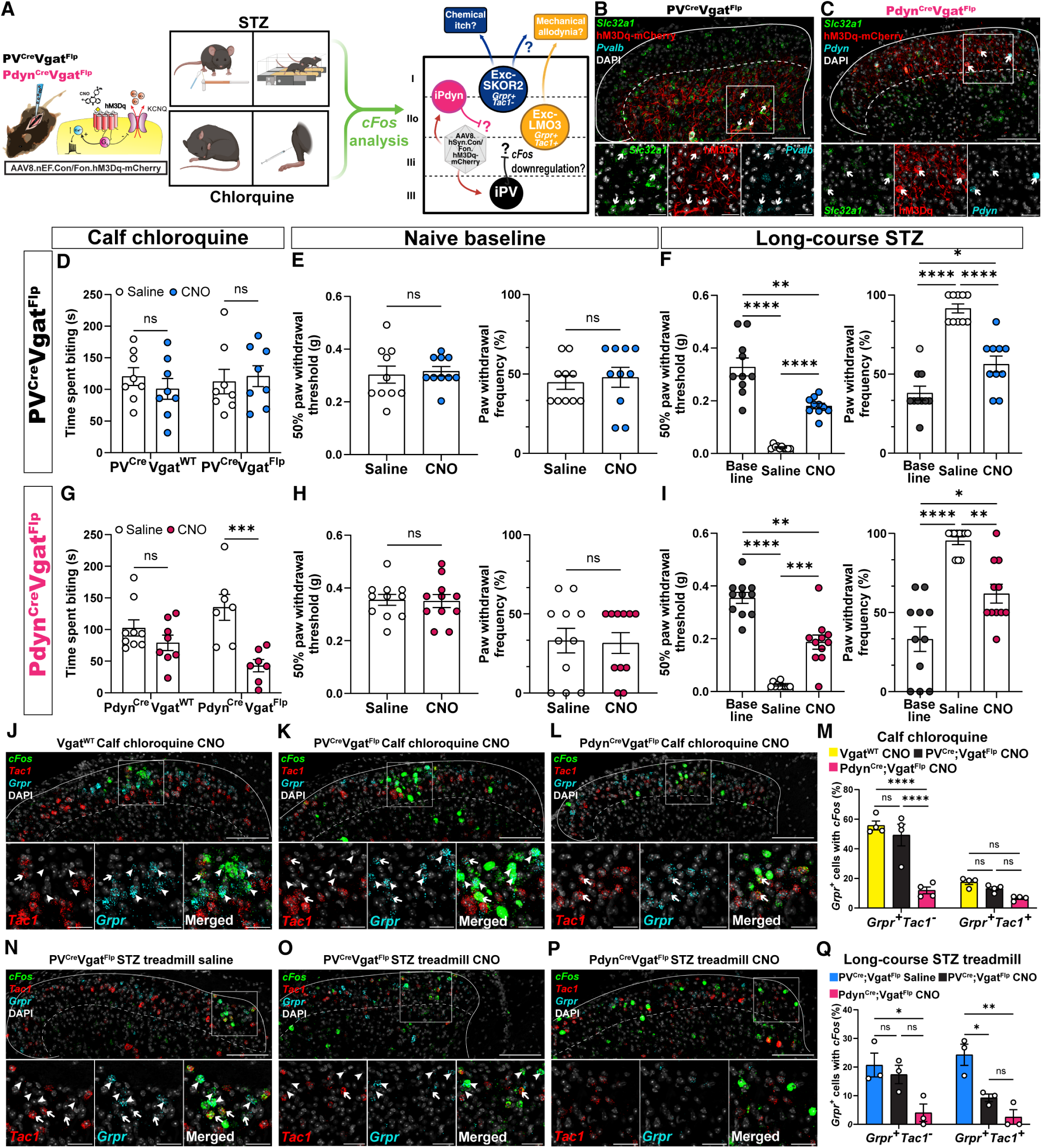
Inhibitory PV and PDYN interneurons differentially regulate functional output of *Grpr* neuron subpopulations. **(A)** Experimental strategy to activate distinct inhibitory interneuron populations, quantify *cFos* expression in Exc-SKOR2 (Grpr⁺Tac1⁻) and Exc-LMO3 (Grpr⁺Tac1⁺) subtypes, and correlate these changes with chemical itch and mechanical allodynia behaviours. **(B and C)** Lumbar dorsal horn sections showing Cre- and Flp-dependent hM3D-mCherry expression (red), *Slc32a1* (green), and (B) *parvalbumin* (*Pvalb*, cyan) transcripts in a PV^Cre^Vgat^Flp^ mouse or (C) *prodynorphin* (*Pdyn*, cyan) transcripts in a Pdyn^Cre^Vgat^Flp^ mouse. Arrows depict cells co-expressing hM3D-mCherry, *Slc32a1*, and (B) *Pvalb or* (C) *Pdyn*. **(D)** Itch-like behaviour after intradermal chloroquine, with saline or CNO, in PV^Cre^Vgat^WT^ (n=8) and PV^Cre^Vgat^Flp^ (n=8) mice. **(E and F)** 50% paw withdrawal threshold (von Frey test, left) and paw withdrawal frequency (cotton swab test, right) at baseline, with saline, or CNO, in (E) naïve (n=10) and (F) long-course STZ (n=10) PV^Cre^Vgat^Flp^ mice. **(G)** Itch-like behaviour after intradermal chloroquine, with saline or CNO, in Pdyn^Cre^Vgat^WT^ (n=9 saline, n=8 CNO) and Pdyn^Cre^Vgat^Flp^ (n=7) mice. **(H and I)** 50% paw withdrawal threshold (von Frey test, left) and paw withdrawal frequency (cotton swab test, right) at baseline, with saline, or CNO, in (H) naïve (n=11) and (I) long-course STZ (n=11) Pdyn^Cre^Vgat^Flp^ mice. (**J to L)** Lumbar dorsal horn sections showing *cFos* (green), *Tac1* (red), and *Grpr* (cyan) transcripts after intradermal chloroquine injection with CNO treatment in (J) Vgat^WT^, (K) PV^Cre^Vgat^Flp^, and (L) Pdyn^Cre^Vgat^Flp^ mice. Arrowheads indicate *cFos* expression in *Grpr^+^Tac1^-^* cells. Arrows indicate *cFos* expression in *Grpr^+^Tac1^+^*cells. **(M)** Percentage of *Grpr^+^*cell subpopulations expressing *cFos* in Vgat^WT^ (n=4), PV^Cre^Vgat^Flp^ (n=4), and Pdyn^Cre^Vgat^Flp^ (n=4) mice after calf chloroquine injection with CNO treatment. **(N to P)** Lumbar dorsal horn sections showing *cFos* (green), *Tac1* (red), and *Grpr* (cyan) transcripts after treadmill stimulation in long-course STZ mice, with saline in (N) PV^Cre^Vgat^Flp^ or CNO in (O) PV^Cre^Vgat^Flp^, and (P) Pdyn^Cre^Vgat^Flp^ mice. Arrowheads indicate *cFos* expression in *Grpr^+^Tac1^-^*cells. Arrows indicate *cFos* expression in *Grpr^+^Tac1^+^* cells. **(Q)** Percentage of *Grpr^+^*cell subpopulations expressing *cFos* after long-course STZ treadmill stimulation in PV^Cre^Vgat^Flp^ (saline n=3, CNO n=3) and Pdyn^Cre^Vgat^Flp^ (n=3) mice. Data are mean ± s.e.m. Dashed lines in representative figures (B, C, J to K, and N to P) delineate lamina II/III border. Scale bars: 100 μm (main images), 30 μm (insets). Statistical analysis was performed with two-tailed paired t-test (E and H), two-way ANOVA with Bonferroni’s post-hoc test (D, G, M, and Q), and repeated-measures one-way ANOVA with Bonferroni’s post-hoc test (F and I). *P < 0.05, **P < 0.01, ***P < 0.001, ****P < 0.0001.

For these experiments, we used an intersectional chemogenetic strategy to activate only iPV or iPDYN interneurons (Fig. 3A). Homozygous PV^Cre^ or Pdyn^Cre^ mice were crossed with heterozygous Vgat^Flp^ mice and then intraspinally injected with an AAV vector encoding the Cre- and Flp-dependent excitatory hM3Dq (iPV^hM3Dq^ or iPDYN^hM3Dq^) (Fig. 3, B and C) (*47, 55*). For controls, we tested Cre^+^ littermates negative for Vgat^Flp^ injected with the same AAV vector. This also controlled for Flp-independent expression of the DREADD. No DREADD was detected in these mice (data not shown). In line with previous reports (*41*), iPV^hM3Gq^ mice exhibited similar chloroquine-induced itch-like behaviour following systemic injection of either CNO (5 mg kg^-1^, i.p.) or saline (i.p.) (Fig. 3D). With respect to *cFos* expression by the two *Grpr^+^* subpopulations in these mice, intradermal chloroquine selectively increased *cFos* in the *Grpr^+^Tac1^-^* neurons, and activation of iPV neurons with CNO had no effect on this upregulation, consistent with the behavioural results (Fig. 3, J, K, and M). In contrast to chemical itch, injection of CNO into iPV^hM3Gq^ mice significantly alleviated both punctate and dynamic mechanical allodynia induced by the long-course STZ model without affecting baseline mechanical thresholds (Fig. 3, E and F). Subsequent innocuous mechanical hindpaw stimulation (treadmill) increased the number of *cFos*^+^ neurons in both *Grpr^+^* subpopulations, as demonstrated earlier, but interestingly, only in the *Grpr^+^Tac1^+^* neuron subtype did CNO-mediated activation of iPV neurons significantly attenuate *cFos* expression (Fig. 3, N, O, and Q).

We then directly tested whether the iPDYN neurons modulate chemical itch and/or mechanical allodynia and whether they do so through one or both *Grpr^+^* subpopulations. In stark contrast to the iPV neurons, activating iPDYN neurons by injection of CNO markedly attenuated chloroquine induced itch-like behaviour (Fig. 3G). As hypothesized, *cFos* upregulation in the *Grpr^+^Tac1^-^*subpopulation due to chloroquine injection was suppressed when iPDYN neurons were activated (Fig. 3, J, L, and M). With respect to mechanical allodynia, like the iPV neurons, activation of iPDYN neurons significantly alleviated both punctate and dynamic mechanical allodynia in the long-course STZ mice without affecting baseline mechanical thresholds (Fig. 3, H and I). In terms of the *cFos* expression, activation of iPDYN neurons with CNO significantly suppressed *cFos* upregulation in both *Grpr^+^* subpopulations (Fig. 3, N, P, and Q). Together, these results suggest a model in which *Grpr^+^Tac1^-^* neurons are important for chemical itch and are inhibited by iPDYN neurons, while *Grpr^+^Tac1^+^*neurons are important for mechanical allodynia and are inhibited by both iPV and iPDYN neurons. That said, while *cFos* analysis provides a correlate of neuronal activity, it does not establish a causal link between the activity of either *Grpr^+^* subpopulation and the behavioural responses to chemical itch or mechanical allodynia.

### Distinct *Grpr^+^* neuron subtypes are necessary for itch and allodynia

To directly test the role of the *Grpr^+^* subpopulations in chemical itch and mechanical allodynia, we employed two complementary approaches that allowed us to capture and manipulate the individual *Grpr^+^* subpopulations (Fig. 4, A and B). The strategy was based on the following observations: First, mechanical allodynia induced by inflammatory or neuropathic injury is attenuated by chemogenetic inhibition of *Grpr* neurons (Fig. 1) but not by ablation of *Grpr* neurons using intrathecally delivered bombesin-saporin (*46*). Second, the *Grpr^+^Tac1^-^*subpopulation preferentially upregulates *cFos* in response to chloroquine (Fig. 2N). And third, not all *Grpr^+^* neurons respond to GRP, the endogenous ligand for the GRPR, or to peripherally administered pruritogen (Fig. 2N) (*14, 49*). Taken together, we hypothesized that bombesin-saporin preferentially ablates *Grpr^+^* neurons that are most responsive to bombesin (i.e. *Grpr*^+^*Tac1*^-^) and this subset of neurons is important for itch (*14, 49*), while the remaining *Grpr^+^* neurons are important for mechanical allodynia (i.e. *Grpr^+^Tac1*^+^). To test this hypothesis, we expressed the hM4Di in *Grpr^+^* neurons by injecting AAV8.hSyn.DIO.hM4Di-mCherry into the lumbar dorsal horn of homozygous Grpr^Cre^ mice. Next, we injected bombesin-saporin intrathecally to selectively ablate *Grpr^+^* neurons. To confirm the selectivity of this ablation, we combined FISH and IF in spinal cord sections from Grpr^hM4Di^ mice treated with either blank-saporin (300 ng, i.t.) or bombesin-saporin (300 ng, i.t.) (Fig. 4, C and D). As expected, bombesin-saporin dramatically reduced the average number of hM4Di^+^ cells in the dorsal horn (Fig. 4E). While approximately 17% of the hM4Di^+^ cells remained after bombesin-saporin treatment (similar to the percent reported previously (*46*)), the composition of this remaining population was significantly skewed to the *Grpr^+^Tac1^+^*subpopulation (Fig. 4F). These results confirm that bombesin-saporin, which selectively attenuates chemical itch behaviour (*46*), preferentially ablates the *Grpr^+^Tac1^-^* neurons, leaving the *Grpr^+^Tac1^+^*neurons intact to be subsequently manipulated using preinjected hM4Di.

**Fig. 4:**
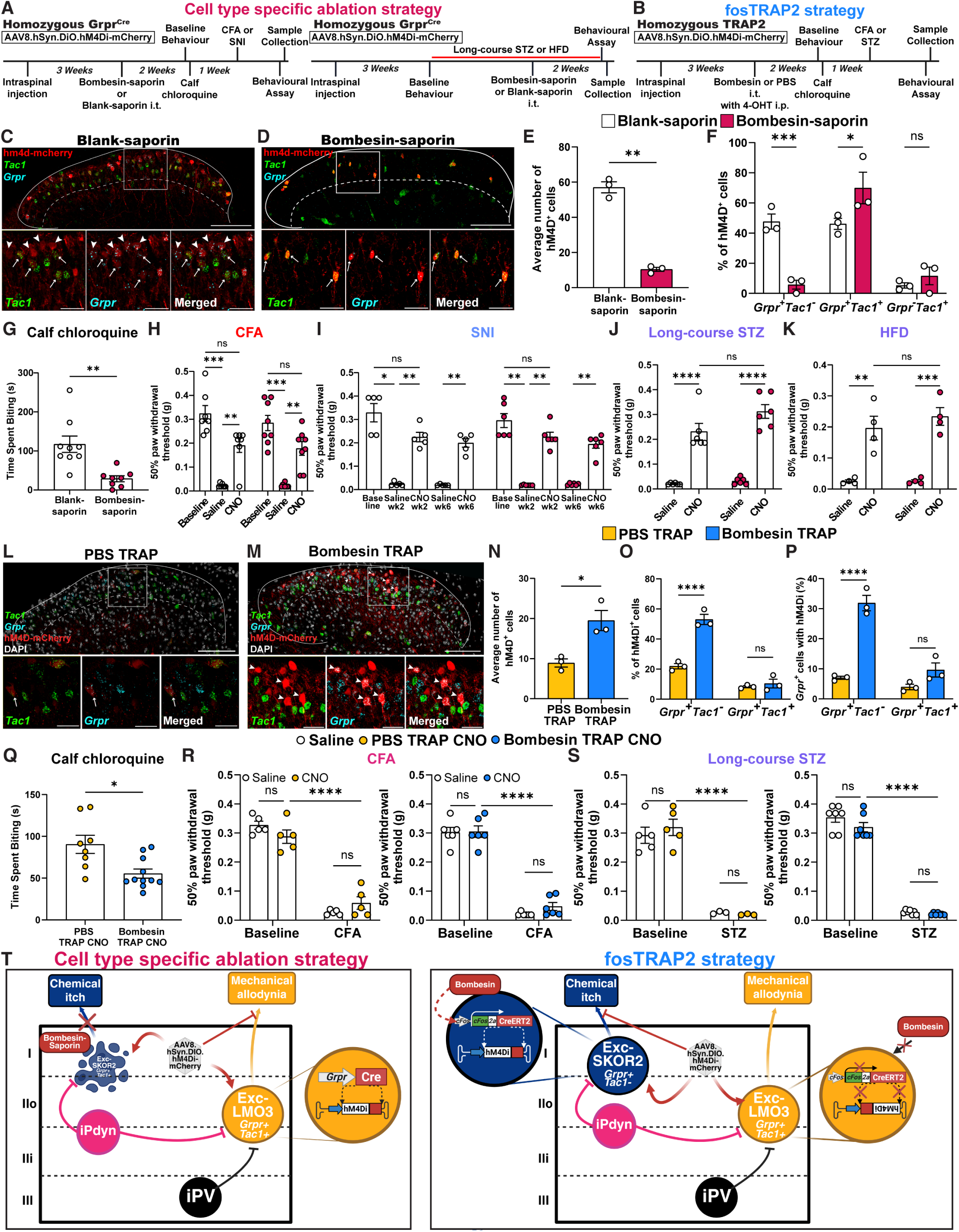
Functional analysis of *Grpr^+^* neuron subtypes in chemical itch and mechanical allodynia. **(A)** Experimental strategy for ablation of bombesin-responsive *Grpr^+^* neurons and chemogenetic silencing of remaining *Grpr^+^* neurons in homozygous Grpr^Cre^ mice. **(B)** Experimental design to selectively silence bombesin-responsive *Grpr^+^* neurons in homozygous TRAP2 mice. **(C and D)** Lumbar dorsal horn sections showing hM4Di-mCherry (red), *Tac1* (green), and *Grpr* (cyan) transcripts after (C) blank- or (D) bombesin-saporin treatment in Grpr^Cre^ mice. **(E)** Average number of hM4Di^+^ cells in blank-(n=3) and bombesin-saporin (n=3) treated Grpr^Cre^ mice. **(F)** Percentage of hM4Di^+^ cells expressing *Grpr^+^Tac1^-^*, *Grpr^+^Tac1^+^*, or *Grpr^-^Tac1^+^* in blank- (white, n=3) and bombesin-saporin (magenta, n=3) treated Grpr^Cre^ mice. **(G)** Itch-like behaviour after intradermal chloroquine injection in blank- (n=9) or bombesin-saporin (n=8) treated Grpr^Cre^ mice. **(H to K)** 50% paw withdrawal threshold with saline or CNO treatment in blank- (white) or bombesin-saporin (magenta) treated Grpr^Cre^ mice: (H) CFA [n=9, 8; same mice as (G)], (I) SNI (n=5, 6), (J) long-course STZ (n=6, 6), and (K) HFD (same mice from Fig. 1i were used; n=4, 4). **(L and M)** Lumbar dorsal horn sections from (L) PBS and (M) bombesin TRAP mice showing hM4Di-mCherry (red), *Tac1* (green), and *Grpr* (cyan) transcript. **(N)** Average number of hM4Di^+^ cells in PBS (n=3) and bombesin TRAP (n=3) mice. **(O)** Percentage of hM4Di^+^ cells expressing *Grpr^+^Tac1^-^* or *Grpr^+^Tac1^+^* transcripts in PBS (blue, n=3) and bombesin TRAP (magenta, n=3) mice. **(P)** Percentage of *Grpr^+^Tac1^-^* and *Grpr^+^Tac1^+^*cells with hM4Di in PBS (blue) and bombesin TRAP (magenta) mice (n=3, 3). **(Q)** Itch-like behaviour after intradermal chloroquine injection with CNO in PBS (n=8) or bombesin TRAP (n=11) mice. **(R and S)** 50% paw withdrawal threshold with saline or CNO in (R) CFA (PBS TRAP n=5, bombesin TRAP n=6) and (S) long-course STZ (PBS TRAP n=5 saline, n=3 CNO; bombesin TRAP n=7) TRAP2 mice. Mice from (Q) were randomly distributed for (R and S). **(T)** Summary of findings from cell type–specific ablation (left) and fosTRAP2 approaches, depicting the roles of Exc-SKOR2 (Grpr⁺Tac1⁻) and Exc-LMO3 (Grpr⁺Tac1⁺) neuron subtypes in mediating chemical itch and mechanical allodynia, respectively. Data are mean ± s.e.m. Dashed lines in representative figures (C, D, L, and M) delineate lamina II/III border; arrowheads indicate hM4Di expression in *Grpr^+^Tac1^-^* cells and arrows indicate hM4Di expression in *Grpr^+^Tac1^+^* cells. Scale bars: 100 μm (main images), 30 μm (insets). Statistical analysis was performed with unpaired two-tailed Welch’s t-test (E, G, N, and Q), and repeated-measures two-way ANOVA with Bonferroni’s post-hoc test (H and I), two-way ANOVA with Bonferroni’s post-hoc test (F, J, K, O, P, R, and S). *P < 0.05, **P < 0.01, ***P < 0.001, ****P < 0.0001.

As a second approach, we selectively captured bombesin-responsive *Grpr^+^* neurons with the chemogenetic receptor hM4Di using tamoxifen-dependent TRAP2 mice (Fig. 4B). The TRAP2 mice use the *cFos* promoter to drive the expression of CreERT2 in response to neuronal activity (*56*). The tamoxifen allows the Cre to be temporarily active on the order of hours (*56*). We injected AAV8.hSyn.DIO.hM4Di-mCherry in the lumbar dorsal horn of the mice. Three weeks later, we intrathecally injected bombesin (0.1 nmol) together with an i.p. injection of 4-hydroxytamoxifen (4-OHT, 50 mg kg^-1^). As a control, we administered PBS (i.t) rather than bombesin together with 4-OHT. To confirm the identity of TRAP^hM4Di^ neurons, we performed FISH and IF in spinal cord sections from PBS TRAP^hM4Di^ and bombesin TRAP^hM4Di^ mice (Fig. 4, L and M). Crucially, bombesin TRAP^hM4Di^ mice showed a differential enrichment of the two major *Grpr^+^*subpopulations compared to PBS TRAP^hM4Di^ (Fig. 4, N to P). The population of bombesin TRAP^hM4Di^ neurons was highly enriched for *Grpr^+^Tac1^-^* neurons (∼53%) compared to PBS TRAP^hM4Di^ (∼19%), with a significantly lower percentage of *Grpr^+^Tac1^+^* neurons (∼10%) which was comparable to the PBS TRAP^hM4Di^ (∼8%) (Fig. 4O). Furthermore, examining the proportion of each *Grpr^+^* neuron subtype captured by the TRAP2 strategy revealed that approximately 32% of all *Grpr^+^Tac1^-^* neurons expressed TRAP^hM4Di^ using bombesin, compared to only ∼7% with PBS (Fig. 4P). Conversely, only ∼9% of *Grpr^+^Tac1^+^*neurons expressed TRAP^hM4Di^ following bombesin injection, a level comparable to the ∼4% observed with PBS (Fig. 4P). These data further reinforce that intrathecal bombesin preferentially activates the *Grpr^+^Tac1^-^*neuron subtype and demonstrate that the TRAP2 approach allows for the selective manipulation of the *Grpr^+^Tac1^-^* subpopulation.

Having established the identities of the captured *Grpr^+^*subpopulations in each approach, we next tested their behavioural contributions to chemical itch and mechanical allodynia induced by multiple injury types. In the case of the bombesin-saporin treated Grpr^hM4Di^ mice, as expected, we observed a significant decrease in chloroquine induced itch-like behaviour (Fig. 4G). Similarly, in the case of the bombesin TRAP ^hM4Di^ mice, administration of CNO (5 mg kg^-1^, i.p.) significantly decreased chloroquine induced itch-like behaviour compared to PBS TRAP^hM4Di^ mice administered CNO (Fig. 4Q). We next tested whether the *Grpr^+^* neurons remaining in the bombesin-saporin treated Grpr^hM4Di^ mice, predominantly belonging to the *Grpr^+^Tac1^+^* subpopulation, are important for mechanical allodynia. Grpr^hM4Di^ mice were first injected with either blank-saporin (300 ng, i.t.) or bombesin-saporin (300 ng, i.t.) and two weeks later, were tested for baseline mechanical sensitivity with von Frey and cotton swab assays, comparing the administration of CNO (5 mg kg^-1^, i.p.) to saline (i.p.). The Grpr^hM4Di^ mice subjected to intrathecal bombesin-saporin or blank-saporin injection showed similar baseline mechanical sensitivity (Fig. 4, H and I). We then tested the mechanical thresholds after induction of the CFA, SNI and long-course STZ models as well as in a second diabetic polyneuropathic pain model, the HFD. For CFA model, the same mice from Fig. 4G were used to directly compare itch and mechanical allodynia behaviours; for diabetic polyneuropathic pain models, bombesin-saporin was administered after the onset of diabetes to minimize potential ablation induced plasticity in the dorsal horn (Fig. 4A). As expected, the blank-saporin treated Grpr^hM4Di^ mice showed significantly attenuated punctate mechanical allodynia in all four models following the administration of CNO compared to systemic saline injection (Fig. 4, H to K). In line with previous findings (*46*), mice treated with bombesin-saporin, which preferentially ablates *Grpr^+^Tac1^-^*neurons, showed normal mechanical allodynia induced by the four injury models (Fig. 4, H to K). However, systemic administration of CNO in the bombesin-saporin treated mice resulted in a robust attenuation of punctate mechanical allodynia across all four injury models compared to systemic administration of saline (i.p.) (Fig. 4, H to K). These results demonstrate that *Grpr^+^Tac1^+^* neurons are necessary and sufficient for mechanical allodynia, while the *Grpr^+^Tac1^-^*neurons are dispensable (Fig. 4T).

We next tested the bombesin TRAP^hM4Di^ mice, which predominantly express hM4Di in the *Grpr^+^Tac1^-^* subpopulation. Here, the same mice from Fig. 4Q were randomly distributed for CFA and long-course STZ models, with extra mice that were not tested for chemical itch. Systemic administration of CNO (5mg kg^-1^, i.p.) had no effect on mechanical allodynia compared to saline (i.p.) in either model (Fig. 4, R and S). As expected, this was also the case for the PBS TRAP^hM4Di^ control mice administered with the same treatments. Taken together, these results demonstrate that the *Grpr^+^Tac1^-^*subpopulation is important for chemical itch and dispensable for mechanical allodynia, while the *Grpr^+^Tac1^+^* neuron subpopulation is important for mechanical allodynia across diverse injury types (Fig. 4T).

### Genomic enhancers enable selective targeting of *Grpr^+^* neuron subtypes

While our previous experiments established the necessity of distinct *Grpr^+^* neuron subtypes in chemical itch and mechanical allodynia, they relied on ablation, which can induce compensatory changes in the dorsal horn circuity, and on indirect methods, i.e. fosTRAP2. To achieve direct, selective, recombinase-independent genetic access to these critical neuronal subpopulations, we utilized genomic enhancers identified with a novel regulatory element screening pipeline (*57*). This enhancer screening approach combines cross-species single-nucleus transposase-accessible chromatin sequencing (snATAC-seq) data from macaque and mouse spinal dorsal horn with machine learning (ML) models to identify and prioritize enhancer sequences predicted to drive cell type specific-gene expression across species (*26, 30, 57*). The strategy represents a significant departure from previous methods and offers the potential for precise genetic targeting in WT animals and facilitating translation across species. First, we identified candidate enhancers using models trained on macaque dorsal horn snATAC-seq data and designed to be specific for a cross-species defined neuron cluster (*57*). This led to the identification of a candidate regulatory element, Excit-1, which our computational models predicted would drive transgene expression in the Exc-LMO3 cluster in mice (*57*). As previously shown, *Grpr^+^Tac1^+^* neurons are assigned almost exclusively to this cluster (Fig. 2, E and F). In a second training of the models, we leveraged both macaque and mouse dorsal horn snATAC-seq data to generate cross-species candidate enhancers for the neuron subtypes. From this, we identified a series of candidate enhancers that were predicted to drive transgene expression in the Exc-SKOR2 subtype (*57*), the cluster that is enriched in *Grpr^+^Tac1*^-^ neurons (Fig. 2, E and F).

Next, we examined the expression pattern of the candidate enhancer for the LMO3 subtype (Excit-1) and one of the candidate enhancers for Exc-SKOR2 (SKOR2.103). AAV8 viruses were constructed with each enhancer driving expression of hM4Di-2a-EGFP and were intraspinally injected into the lumbar dorsal horns of WT mice (Fig. 5, A and B). Expression analysis performed three weeks later showed the Excit-1 enhancer virus conferred robust expression, while the SKOR2.103 driven expression was dimmer, requiring the addition of a minimal beta-globin (mBG) promoter. Analyzing the distribution of EGFP within the two major *Grpr*^+^ subpopulations with FISH and IF, expression of the Excit-1-driven hM4Di-2a-EGFP (Excit-1^hM4Di^) was highly selective for *Grpr^+^Tac1^+^*neurons, with ∼40% of hM4Di^+^ cells overlapping with *Grpr^+^Tac1^+^*, and only ∼9% with *Grpr^+^Tac1^-^* (Fig. 5C). The SKOR2.103mBG-driven expression (SKOR2.103^hM4Di^), however, while overlapping with both *Grpr^+^* subpopulations, showed greater enrichment in *Grpr^+^Tac1^-^*compared to the Excit-1^hM4Di^ virus (Fig. 5C). With respect to the percentage of each subpopulation transduced by the two viruses, Excit-1^hM4Di^ captured a significant percentage of the *Grpr^+^Tac1^+^* subpopulation (∼55%) but few of the *Grpr^+^Tac1^-^*neurons (∼11%), compared to SKOR2.103mBG ^hM4Di^ which captured a significant percentage of both populations (∼49% of *Grpr^+^Tac1^-^*and ∼70% of *Grpr^+^Tac1^+^*neurons) (Fig. 5C). No EGFP^+^ neurons were detected in the L3-L5 dorsal root ganglia (DRG) neurons or brains of mice injected with either enhancer virus (fig. S6A and not shown). These results validate the Excit-1 element as highly selective tool for targeting *Grpr^+^Tac1^+^*neurons, while SKOR2.103mBG targets *Grpr^+^* neurons more broadly.

**Fig. 5:**
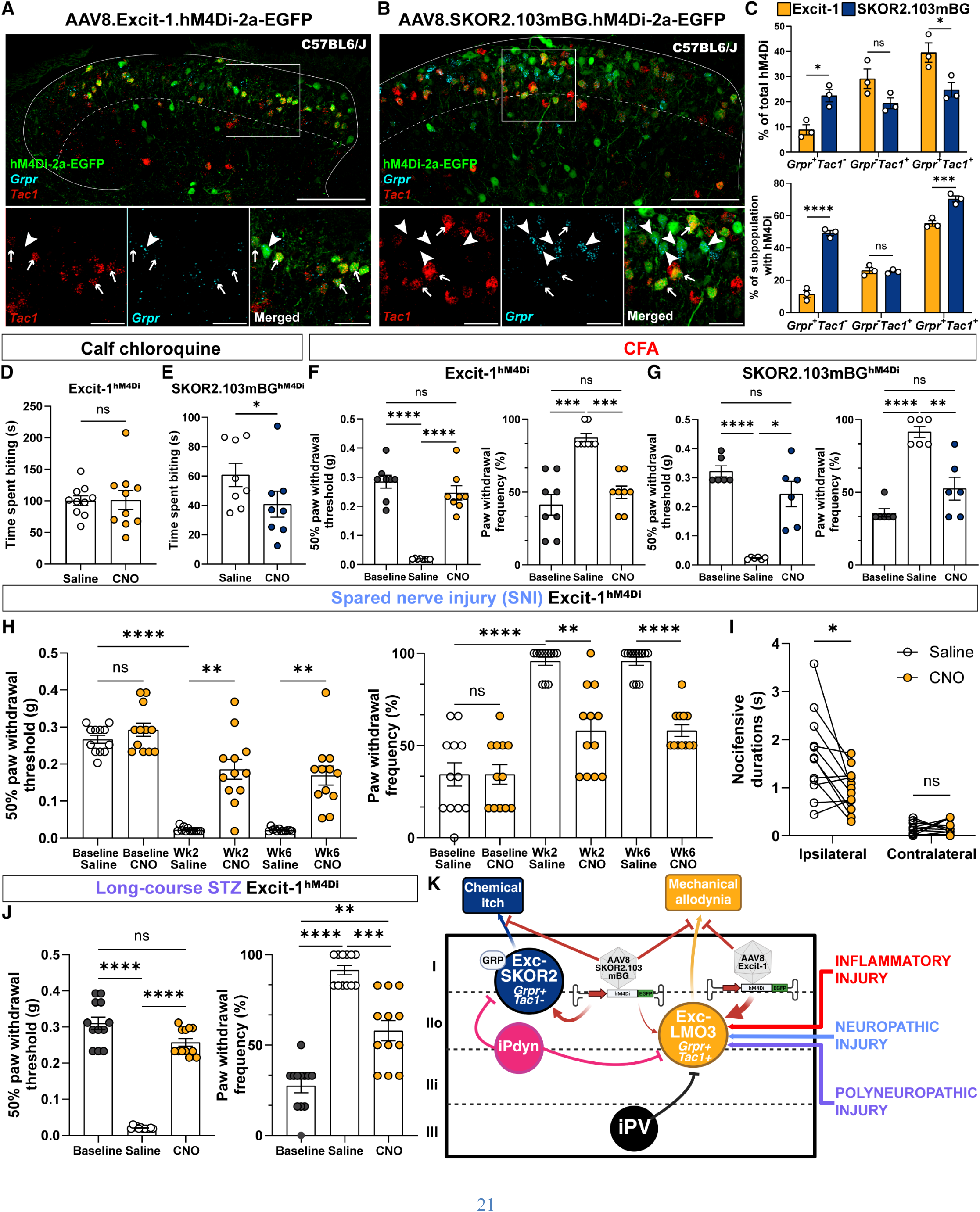
Computationally identified genomic enhancers target *Grpr^+^* neuron subtypes to differentially modulate itch and mechanical allodynia. (A and B) Lumbar dorsal horn section showing hM4Di-2a-EGFP (green), *Tac1* (red), and *Grpr* (cyan) expression following intraspinal delivery of AAV8.Excit-1.hM4Di-2a-EGFP (A) or AAV8.SKOR2.103mBG.hM4Di-2a-EGFP (B) in wild type mice. Dashed lines delineate lamina II/III border. Arrowheads indicate *Grpr^+^Tac1^-^* cells with hM4Di and arrows indicate *Grpr^+^Tac1^+^* cells with hM4Di. Scale bars: 100 μm (main images), 30 μm (insets). **(C)** Quantification of enhancer selectivity and coverage. The top panel shows the distribution of all enhancer-driven hM4Di^+^ cells across *Grpr^+^Tac1^-^*, *Grpr^-^Tac1^+^*, and *Grpr^+^Tac1^+^* subtypes for Excit-1 and SKOR2.103mBG enhancers. The bottom panel shows the percentage of cells within each endogenous subtype that express hM4Di driven by either enhancer (n=3 mice per enhancer). **(D and E)** Itch-like behaviour after intradermal chloroquine injection with saline or CNO treatment in mice expressing Excit-1^hM4Di^ or SKOR2.103mBG^hM4Di^ (n=10 and 8). **(F to H),** 50% paw withdrawal threshold (von Frey test, left) and frequency (cotton swab test, right) with saline or CNO in (F and G) CFA (n=8 and 6), and (H) SNI (n=12), and (J) long-course STZ (n=12) mice with Excit-1 (F, H, J) or SKOR2.103mBG (G) element driven spinal hM4Di expression. The same cohort of mice used in (D) was randomly assigned to experiments shown in (F) and (H), while mice from (E) were used for experiment (G). **(I)** Duration of nocifensive behaviours (shaking, lifting, and licking) in SNI mice (>6 weeks post-injury) expressing Excit-1^hM4Di^ with saline or CNO (n=12; same mice as (H)). **(J)** 50% paw withdrawal threshold (von Frey test, left) and frequency (cotton swab test, right) with saline or CNO in long-course STZ (n=12) mice intraspinally injected with AAV8.Excit-1.hM4Di-2a-EGFP after the onset of STZ induced diabetes. **(K)** Summary of findings in this study depicting a molecular and spinal circuit basis for the functional segregation of itch and pain. Data are mean ± s.e.m. Statistical analysis was performed with unpaired two-tailed Welch’s t-test (D), paired two-tailed t-test (E), repeated-measures one-way ANOVA with Bonferroni’s post-hoc test (F to H and J), and repeated-measures two-way ANOVA with Bonferroni’s post-hoc test (I). *P < 0.05, **P < 0.01, ***P < 0.001, ****P < 0.0001.

Next, we tested the functional role of Excit-1^hM4Di^ and SKOR2.103^hM4Di^ captured neurons in chemical itch and mechanical allodynia behaviour. Administration of CNO (5mg kg^-1^, i.p.) to Excit-1^hM4Di^ injected mice showed no effect on chloroquine-induced itch-like behaviour (Fig. 5D), while silencing SKOR2.103^hM4Di^ expressing neurons significantly attenuated itch-like behaviour (Fig. 5E). As a negative control, we generated a virus using only mBG to drive expression of the inhibitory chemogenetic receptor (fig. S7A) (mBG^hM4Di^). Chemogenetic silencing of mBG targeted neurons did not affect chemical itch behaviour (fig. S7B). Together, these findings reinforce the role of the *Grpr^+^Tac1^-^* (i.e. Exc-SKOR) subpopulation in mediating chemical itch. With respect to mechanical allodynia, silencing Excit-1^hM4Di^ expressing neurons, in the same mice tested in Fig. 5D, significantly attenuated punctate and dynamic mechanical allodynia in the CFA model, consistent with its significant expression by *Grpr^+^Tac1^+^*neurons (Fig. 5F). Similarly, silencing the SKOR2.103^hM4Di^ expressing neurons, which likewise transduce a substantial portion of *Grpr^+^Tac1^+^* neurons, also significantly attenuated punctate and dynamic mechanical allodynia in the CFA model (Fig. 5G). By contrast, silencing mBG^hM4Di^ targeted neurons had no effect on punctate or dynamic mechanical allodynia in the CFA model (fig. S7C). Furthermore, in the SNI model, silencing Excit-1^hM4Di^ neurons not only reduced both punctate and dynamic mechanical allodynia, without affecting baseline mechanical thresholds, but also significantly attenuated the duration of nocifensive behaviours, such as licking, shaking, and lifting, which are associated with sustained pain (Fig. 5I) (*58, 59*). Lastly, to assess the therapeutic potential of this approach in a more clinically relevant setting, we injected AAV8.Excit-1.hM4Di-2a-EGFP intraspinally 4 weeks after the onset STZ-induced diabetes. CNO administration two weeks later produced a robust recovery of both punctate and dynamic mechanical allodynia (Fig. 5J). Novel genomic enhancers, such as these, provide powerful tools for direct and selective targeting of dorsal horn neuron subtypes, enabling the investigation of their specific contributions to somatosensory processing and allowing for the implementation of new cell type targeted, mechanism-based therapeutic strategies for chronic pain.

## Discussion

Deciphering the neural codes for distinct somatosensory modalities like itch and pain within the intricate circuitry of the spinal dorsal horn remains a central goal in neurobiology. For decades, the field has been challenged by the inability to parse spinal neurons into their fundamental functional units and often conflating neuronal responses to stimuli with functional necessity. These challenges are amplified by the profound cellular heterogeneity of the spinal dorsal horn and complexities introduced by chronic pain states, where maladaptive plasticity can alter sensory processing (*2, 3*). Our investigation began by exploring the circuits underlying mechanical allodynia in polyneuropathic injury models, leading us unexpectedly to re-examine the role of *Grpr^+^* neurons. Remarkably, by moving beyond a single gene marker, we discover that closely related, but transcriptomically distinct, interneuron subtypes can mediate qualitatively different sensations, highlighting the power of integrating transcriptomics with functional manipulations to resolve complex coding logic within somatosensory circuits.

Our initial finding that mechanical allodynia in polyneuropathic injury models recruits a unique neuronal ensemble, distinct from those mediating inflammatory or neuropathic injuries, was surprising given polyneuropathic injuries are a subtype of neuropathic injury (fig. S1) (*33*). The key pathways involving VGLUT3, CCK, PKCγ, and CR neurons previously implicated in mechanical allodynia (*31, 32, 60*), were dispensable for polyneuropathic states. This divergence underscores the observation that spinal circuits that mediate mechanical allodynia are heterogeneous; with different aetiologies recruiting distinct spinal pathways, and thus requiring more nuanced, mechanism-based therapeutic strategies.

While *Grpr^+^* neurons possess the anatomical characteristics of vertical cells, implicated in mechanical allodynia (*2, 3, 14, 37, 39*), prior work suggested they were exclusively dedicated to itch and dispensable for pain (*44–46, 61*). Our analysis of the function of these neurons using broad chemogenetic silencing, revealed a critical role in mechanical allodynia across all tested injury types, including polyneuropathic injuries (Fig. 1). This created a clear paradox: how could neurons dedicated to itch be essential for diverse forms of mechanical allodynia? We resolved this paradox by demonstrating that *Grpr^+^* neurons are not homogeneous. Single cell and spatial transcriptomic analyses, FISH, and mapping onto the species-conserved atlas of dorsal horn neuron subtypes (*26*), revealed two major *Grpr^+^* subpopulations defined by the differential expression of *Tac1*. Notably, we showed that the *Grpr^+^Tac1^-^* neurons are important for itch and trended towards higher *Grpr* transcript levels. These higher levels potentially explain their responsiveness to GRPR ligands (*14, 49*), which are known mediators of itch in the spinal dorsal horn (*40, 45, 61*). Concomitantly, we reinforce the previous findings that GRPR and its ligands in the dorsal horn are dispensable for mechanical allodynia (*45, 61*).

Multiple lines of evidence provided here support a functional dichotomy for the otherwise closely related *Grpr*^+^ subpopulations (Exc-SKOR2 and Exc-LMO3 neuron subtypes). An interesting question pertaining to these findings is how the two dorsal horn neuron subtypes differentially mediate itch and pain signals given the largely polymodal nature of innervating nociceptors, including their response to pruritogens (*6–12*). Our results suggest one key influence may be the differential control by inhibitory interneurons. We found that while activating iPV interneurons had no effect on chloroquine induced itch-like behaviour, activating iPDYN interneurons reduced itch behaviour and correspondingly prevented the upregulation of *cFos* in *Grpr^+^Tac1^-^* neurons (Fig. 3). Although activation of either iPV and iPDYN neurons reduced polyneuropathic injury induced allodynia as well as *cFos* upregulation in the *Grpr^+^Tac1^+^*neurons (Fig. 3), the selective inhibitory modulation of itch suggests distinct inhibitory gating mechanisms shape the modality specific output. It is unclear how these inhibitory interneurons are engaged in a natural environment; however, distinct patterns of primary afferent activation likely play a role (*5*). Indeed, activation of different classes of nociceptors *in vitro* led to increased inhibitory postsynaptic currents (IPSCs) in *Grpr^+^* neurons (*42*). Moreover, the neuropeptidergic milieu of the dorsal horn likely acts in concert with distinct patterns of primary afferent activation to further refine modality coding (*62*). Notably, somatostatin-mediated suppression of iPDYN neurons can disinhibit *Grpr^+^* neuron-mediated itch pathways (*51, 52*), and natriuretic polypeptide b (Nppb) mediated activation of GRP neurons (*63*) can potentiate *Grpr^+^* neuron activity through GRP (*40*), which we demonstrate preferentially activates the *Grpr^+^Tac1^-^*neuron subtype (Fig. 4). We thus propose a model whereby the overall balance of excitation and inhibition in the dorsal horn, shaped by both synaptic inputs and the peptidergic influences, dictates the preferential activation of either the *Grpr^+^Tac1^-^*(itch), or *Grpr^+^Tac1^+^* (allodynia) pathway, allowing the dorsal horn to parse complex, polymodal information into modality specific output (Fig. 5K).

A key methodological advance in this study is our validation and utilization of the computationally designed genomic enhancers, Excit-1 and SKOR2.103mBG. These tools provided unprecedented, selective genetic access to the *Grpr^+^Tac1^+^* (Exc-LMO3) and *Grpr^+^Tac1^-^*(Exc-SKOR2) subpopulations, respectively, allowing us to link each subtype to a distinct behavioural output. Specifically, the ability of Excit-1-mediated silencing to reverse mechanical allodynia across diverse injury states without affecting itch, contrasted by the attenuation of itch by silencing SKOR2.103mBG-targed neurons, provides direct causal evidence for the functional dichotomy of *Grpr^+^*neuron subtypes. Furthermore, the efficacy of activating Excit-1^hM4Di^ in reducing nocifensive behaviours associated with sustained pain in neuropathic pain model and reversing polyneuropathic mechanical allodynia, even when delivered post-injury, underscores its therapeutic potential. Identifying the most robust and specific genomic enhancers for dorsal horn neuron subtypes across species is made possible by efficiently scaling the process with high-throughput screening approaches, such as the novel methods described by Leone *et al.* in this same issue (*57*). Ultimately, these neuron subtype-enriched enhancers will unlock new avenues for detailed functional studies and circuit mapping as well as investigating how neuron subtype-specific activity patterns shape ascending pathways to generate distinct sensory percepts supraspinally.

## Materials and Methods

### Animals

#### Mice

All animals were kept on a standard 12:12 light/dark cycle with food and water provided *ad libitum* and were treated in accordance with protocols approved by the University of Pittsburgh’s Institutional Animal Care and Use Committee (IACUC). Mouse strains obtained from Jackson Laboratories include: C57BL/6J (RRID: IMSR_JAX:000664), CCK^Cre^ (RRID: IMSR_JAX:012706), CR^Cre^ (RRID: IMSR_JAX:010774), Pdyn^Cre^ (RRID: IMSR_JAX:027958), Grpr^Cre^ (RRID: IMSR_JAX:036668), PV^Cre^ (RRID: IMSR_JAX:017320), TRAP2 (RRID: IMSR_JAX:030323), Vgat^Flp^ (RRID:IMSR_JAX:029591), and Lep^Ob/Ob^ (RRID:IMSR_JAX:000632). VGLUT3KO (RRID: IMSR_JAX:016931), VGLUT3^fl/fl^, and VGLUT3^iCre^ (RRID:IMSR_JAX:018147) mice were generated as previously described (*31, 64, 65*). Dr. Carmen Birchmeir provided the Lbx1^Cre^ mice (*66*) and Drs. David Ginty and Victoria Abraira provided the PKCγ^CreERT2^ mice (*67*). Both males and female mice, unless otherwise specified, between 1.5 to 6 months were used in this study.

#### Macaques

Lumbar spinal cord samples for fluorescence *in situ* hybridization (FISH) were obtained from one *Macaca mulatta* (M94-17; Male, 3 years) provided by Dr. David Lewis and one *Macaca fascicularis* (M263-23; Male, 5 years) provided by Dr. William Stauffer at the University of Pittsburgh. All housing and experimental procedures were performed in accordance with the guidelines of the US Department of Agriculture and the National Institutes of Health Guide for the Care and Use of Laboratory Animals and protocols approved by the University of Pittsburgh’s IACUC.

#### Tamoxifen induction for PKCγ^CreERT2^ transgenic mice

To induce Cre recombinase activity in PKCγ^CreERT2/+^ mice, tamoxifen (Sigma-Aldrich, T5648) was prepared in corn oil to a final concentration of 20mg ml^-1^ as previously described (*32*). One week following intraspinal injection of AAV8.Ef1α.DIO.hM4Di-mCherry, mice received intraperitoneal (i.p.) injections of tamoxifen (75mg kg^-1^) once daily for five consecutive days. All subsequent behavioural and histological experiments were performed at least three weeks after the final tamoxifen injection to allow for maximal transgene expression.

#### Intraspinal injection of adeno-associated virus (AAV) vectors

Intraspinal injections were performed as previously described (*31, 68*) with minor modifications. Mice, 21-28 days (Figs. 1,3,4, and 5, D to I) or 3 months old (Fig. 5J), were deeply anesthetized with 4% isoflurane and maintained with 1.5-2% isoflurane throughout the procedure. A midline incision was made to expose the thoracolumbar vertebral column. Without performing a laminectomy, the intervertebral space between T12-T13 (to target lumbar segments L3-L4 (*69*)) was exposed on the left side with blunt dissection. For most experiments, a single injection was performed (Figs. 1, 3, 4 and fig. S1); however, for experiments utilizing regulatory element-driven AAVs, two injections were performed at intervertebral spaces between T12-T13 and T13-L1 (to target lumbar segments L4-L5 (*69*)) to maximize transgene expression (Fig. 5). The vertebral column was stabilized by clamping at L1 using V-notch spikes (Kopf Instruments). After making a small tear of the dura, a glass microelectrode (tip diameter ∼50 µm) was inserted 350-400 µm lateral to the dorsal midline blood vessel to a depth of 200 µm from the dorsal surface using a stereotaxic frame (Stoelting or Kopf Instruments). AAV vectors (specific details in table S1) were infused over a period of 5 minutes using a stereotaxic injector (Stoelting or Drummond Scientific). The micropipette remained in place for at least 3 minutes post-infusion before being slowly withdrawn. The latissimus dorsi muscle was closed with 6-0 nylon sutures, and the skin was closed with 6-0 silk sutures. Unless otherwise stated, behavioural and histological experiments were performed at least 3 weeks post-injection to allow for maximal and stable viral transgene expression.

#### Monosynaptic retrograde rabies tracing

Monosynaptic retrograde rabies tracing was performed as described previously (*70*), with minor modifications. Briefly, AAV1.hSyn.FLEX.splitTVA-EGFP-tTA and AAV1.TREtight.mTagBFP2-B19G were diluted to 8.5e10 vg mL^-1^ and 3.0e12 vg mL^-1^ respectively in Dulbecco’s phosphate-buffered saline (DPBS; Gibco, 14190-144) and mixed in a 1:1 ratio to a final volume of 500 nL. This helper AAV mixture was intraspinally injected into the lumbar dorsal horn of P21 C57BL/6J (wild-type control) or homozygous Grpr^Cre^ mice, as described above. Seven days post-helper AAV injection, EnvA G-deleted rabies virus encoding dsRedXpress (RVΔG-dsRedXpress; 4.18e8 vg mL^-1^, 350 nL) was intraspinally injected into the same location. Spinal cord and ipsilateral L3-L5 dorsal root ganglia (DRGs) were collected 7 days after the rabies injection for immunofluorescence (IF) and FISH analysis.

#### Construction of Excit-1, SKOR2.103mBG, and mBG AAV8 vectors

Following sequences were used to generate AAV8.Excit-1.hM4Di-2a-EGFP, AAV8.SKOR2.103mBG.hM4Di-2a-EGFP, and AAV8.mBG.hM4Di-2a-EGFP.

Excit-1 sequence:

“GGTTGGAACAATCAACACGTCCAGCATGAGATATGGGCCATTCGCCGACCCAAGTCTAGGCTGTCTCAAGGCCCCCAGGCCTGGTGCCATGGGCACCCCACCAGTCCCCAGCCTGAATGTCCACGCAGGGGGTGGAGTGGGCGGGAGGGCTTCCAGAGTGTGCTGGCACGGAGCGGTCCTCACTGCCCCTGCCTGAGCCTCATCAGTGGCAGACATAAACTAACTCGGTTCCTGGGGACCCAGAGGAGCAGCAGGAACTGTTTGGTAGAGAATTATAGCTGTGGGTGAAATTAGATGTACTCTGAGCCCCAAGGCTGAGAGAAATAAAAAGCATTTTAAATGGAGTGTAGAATTCATATTCCTCAGTCACCTTCCAATAAATAAATCTTGTACCATTTTTTTCTGTAAACAGTGGTTACACTCAGCTTCCATTGTCTCCACATCCCATATGCCAATTGGCAGGGAGTAGCAGGGCCGTGCTGGACTGGGAAGGAAAAACAT”

SKOR2.103mBG sequence (small case indicates the mBG sequence):

“ATGTAACTCAGCTGAAGAATCATTATAGGATTTTTTTAAAAACTGGAAACTTCCCCCCTGCCCCCAGCTGTCTACAGAAATAGATTATTTCTTCTAATTTCCTGATTTAGTTATAGTCATAGTGTACATAAAAACAGATTTGATTTTCTTGGAATCAAACATAAAATGAAAGCCAAGGCCTTATCATTGAAACAAGACAGCAGTTTTGCAAAGCTGTGTACATCCTTTGAACTCAGATGACATTGTCTCAGAGGAAAAGATGTTTGATGAGTTTTTTACACTTTAAAGCTCTTCCTAAGGAAAATGAGCTGGGAATCTCCTTTGGTAGAATTGAAGGGGGGTGGATCATTTCGTCTTCAAAAGATGAGGGGAAAAGTGTCATTTTCTATTTCCCTTAAGTATCATTCTTTCAAATACTTGTGTGACAAGAGGTTCAACAGGCTTAGATTTGGGTTTTTAGTTTTCCAGAAATGCATGGAGGACTTAAATGGCCCTGTACAgggctgggcataaaagtcagggcagagccatctattgcttacatttgcttctg”

mBG sequence:

“GGGCTGGGCATAAAAGTCAGGGCAGAGCCATCTATTGCTTACATTTGCTTCTG”

Excit-1, SKOR2.103mBG, mBG gene fragments were made by Twist Bioscience and subcloned into the pAAV.SYN1.HA-hM4Di (Addgene, plasmid# 121538) (*71*) using MluI (New England Biolabs; R3198S) and KpnI (New England Biolabs, R3142S) restriction enzyme sites. HA-hM4Di sequence was replaced by hM4Di-2a-EGFP from pAAV.EF1α.Flex/3’USS.hM4D-2A-EGFP(ATG mut) (Addgene, plasmid # 197892) (*72*) using a PCR kit (New England Biolabs, M0530S) and restriction enzymes KpnI and HindIII (New England Biolabs, R3104S). Whole plasmid sequencing was performed by Plasmidsaurus, and each plasmid was packaged into AAV8 by Neurotools viral vector core (UNC).

#### Chemogenetic activation

To activate PSAM^L141F,Y115F^-GlyR, PSEM^89S^ (30mg kg^-1^; Tocris, 6426), dissolved in sterile saline, was administered i.p. 15 minutes prior to behavioural assessments (fig. S1). For activation of hM4Di (Figs. 1,4,5) or hM3Dq (Fig. 3) DREADDs (*47*), clozapine-N-oxide dihydrochloride (CNO; 5mg kg^-1^; Tocris, 6329), dissolved in sterile saline, was administered i.p. 30 minutes prior to experiments. A crossover experimental design was employed, with identical i.p. saline injections serving as the vehicle control. For all experiments, the experimenter was blinded to the treatment conditions.

### Pain models

#### Oxaliplatin model of chemotherapeutic polyneuropathy

Oxaliplatin model of chemotherapeutic polyneuropathy was induced as previously described (*73*). Oxaliplatin solution was prepared daily in sterile water. VGLUT3^fl/fl^ or VGLUT3^fl/fl^;Lbx1^Cre^ mice received i.p. injections (3mg kg^-1^) daily for 5 days followed by 5 days of no treatment for two cycles (30mg kg^-1^ accumulated dose). Behavioural measurements were performed 20 days after the initial treatment.

#### Long-course streptozotocin model of diabetic polyneuropathy

The long-course streptozotocin (STZ)-induced model of diabetic polyneuropathy was performed as previously described (*74*). Mice received intraperitoneal (i.p.) injections of STZ (60 mg kg^-1^; Sigma-Aldrich, S0130), freshly dissolved in sterile 0.05 M sodium citrate buffer (pH 4.5), once daily for six consecutive days. To confirm hyperglycemia (blood glucose >300 mg dL^-1^), tail vein blood glucose levels were measured daily for two weeks starting 14-16 days after the first STZ injection, and weekly thereafter (fig. S3A). Mice that did not become hyperglycemic within 17-20 days after the initial STZ regimen received an additional STZ injection (60 mg kg^-1^, i.p.). To prevent severe hyperglycemia and death, mice with blood glucose levels exceeding 450 mg dL^-1^ received subcutaneous (s.c.) insulin (1-2 U of 40 U mL^-1^; Merck, 127260). Mechanical withdrawal thresholds were assessed 6 weeks after hyperglycemia confirmation, and treadmill stimulation for cFos analysis was performed 7 weeks after hyperglycemia confirmation.

#### Leptin-deficient (ob/ob) mouse model of diabetic polyneuropathy

Homozygous leptin-deficient mice (Lep^ob/ob^) were acquired from Jackson laboratories. Lep^ob/ob^ mice were crossed to VGLUT3 knockout mice (VGLUT3^-/-^) or VGLUT3 wildtype mice VGLUT3^+/+^ to generate Lep^ob/ob^;VGLUT3^-/-^ or Lep^ob/ob^;VGLUT3^+/+^ respectively. Mechanical thresholds were measured in 5-, 10-, 12-, and 14-week-old mice.

#### High fat diet model of diabetic polyneuropathy

High fat diet (HFD) model of diabetes was induced as previously described (*75*). Male mice were fed a high fat and sucrose content diet (Envigo, TD88137) for 10 weeks. Control mice received a regular diet. Weights were recorded at baseline, 4 weeks, 8 weeks, and 10 weeks (fig. S3B). To confirm diabetes, a glucose tolerance test was performed after 10 weeks: Mice were fasted for 12 hours and their blood glucose levels recorded before they were injected with a 45% d-glucose solution (2000 mg glucose kg^-1^ body weight). Blood glucose was measured 30, 60, and 120 minutes after injection (fig. S3, C and D). Behavioural assessment was performed 10 (Fig. 1K) or 12 weeks (Fig. 4K) after the start of the HFD. Mice in Fig. 4K were from Fig. 1K, which were randomly distributed to two experimental groups.

#### Complete Freund’s Adjuvant model of inflammatory pain

Mice were deeply anesthetized with 4% isoflurane and maintained at 2.5% isoflurane. Glabrous surface of the left hindpaw was injected with 10μL of an emulsion of equal parts Complete Freund’s Adjuvant (CFA) (Sigma, F5881) and sterile saline. Behavioural assays were performed 3-5 days after the CFA injection.

#### Spared nerve injury model of neuropathic pain

Spared nerve injury (SNI) model was performed as described previously (*76, 77*). Mice were deeply anesthetized with 4% isoflurane and maintained at 2% isoflurane. The left hindlimb thigh was shaved and disinfected with 70% ethanol and betadine. A small skin incision was made over the lateral aspect of the thigh, and blunt dissection was performed through the biceps femoris muscle to expose the sciatic nerve and its three terminal branches: the common peroneal, tibial and sural nerves. The common peroneal and tibial nerves were tightly ligated with 6-0 nylon suture and then transected distal to the ligation, with 1-2 mm of the distal nerve stump removed to prevent regeneration. Care was taken to ensure the sural nerve remained intact and undamaged. The muscle layer and the skin were closed with 6-0 nylon sutures. Behavioural assays were performed 1, 2, and 6 weeks after the SNI.

### Behavioural assays

#### Assessment of punctate and dynamic mechanical allodynia

Mice were tested as previously described (*32*). Mice were habituated to opaque Plexiglas chambers on a wire mesh table for at least an hour prior to giving the treatments. For punctate mechanical allodynia assessment, von Frey test was performed using a set of calibrated Semmes-Weinstein monofilaments using the Up-Down method, beginning with a 0.4 g filament (*78*). The 50% paw withdraw threshold (PWT) was determined for each mouse on one or both hind paws. Each filament was gently applied to the plantar surface of the hind paw for 3 s or until a response such as a sharp withdrawal, shaking or licking of the limb was observed. Incidents of rearing or normal ambulation during filament application were not counted. Filaments were applied at five-minute intervals until thresholds were determined. For dynamic mechanical allodynia assessment, the head of a cotton swab was teased and puffed out until it reached approximately three times its original size. The cotton swab was lightly run across the surface of the hind paw from heel to toe. If the animal reacted (lifting, shaking, licking of the paw) a positive response was recorded. A negative response was recorded if the animal showed no such behaviour. The application was repeated 6 times with a 3-minute interval between applications, and a percentage frequency of response determined for each animal. Throughout the experiment, the experimenter remained blinded to treatment conditions, genotypes, and/or AAV vectors injected.

#### Assessment of nocifensive behaviours

Nocifensive behaviours, indicative of sustained pain, was assessed as previously described (*59*). Briefly, the duration of nocifensive responses, such as paw shaking, lifting or hovering, and licking, was evaluated following successful application of a 2 g von Frey filament to the hind paw for 20 seconds. Nocifensive responses were recorded from beneath the testing chamber using a Sony Handycam CX405. Each mouse underwent three successful trials on both the ipsilateral and contralateral paws. Mice received either CNO or vehicle (saline, i.p.) 30 minutes prior to assessment of nocifensive behaviours and a crossover experimental design was employed the next day to assess the effects of CNO treatment within a subject. Videos were subsequently assigned random identification numbers for blinded analysis. Behavioural Observation Research Interactive Software (BORIS) (*79*) was used to measure the duration of nocifensive behaviours. The mean duration was calculated by averaging across the three trials.

#### Assessment of chloroquine induced itch-like behaviour

Itch-like behaviour was assessed following intradermal (i.d.) chloroquine (CQ) injection, as previously described (*80*). Briefly, the calves of experimental mice were shaved one day prior to behavioural testing. On the day of testing, mice were first habituated to the behavioural room for at least 1 hour, followed by a 30-minute habituation period within individual opaque plexiglass chambers placed on a clear plexiglass table. CQ (5 µL of 20 mg mL^-1^ in sterile saline; Sigma-Aldrich, C6628) or vehicle (5 µL sterile saline) was then administered i.d. into the shaved calf with a 30-gauge needle attached to a 25 μL Hamilton syringe (Hamilton company; 80401). Immediately following injection, behaviour was video recorded from underneath the clear plexiglass table for 30 minutes using a Sony Handycam CX405. Videos were subsequently assigned random identification numbers for blinded analysis. The total duration of biting directed towards the injected calf was quantified using BORIS (*79*). All video scoring was performed by experimenters blinded to the treatment conditions and genotypes of the mice. For experiments involving only CQ or saline administration (Fig. 2), C57BL/6J mice were randomly assigned to receive either CQ or vehicle. For experiments involving chemogenetic manipulations, mice received either CNO or vehicle (saline, i.p.) 30 minutes prior to i.d. CQ administration. Unless otherwise specified, a crossover design was employed for these chemogenetic experiments, with at least 3 days between treatments. However, for the experiment depicted in Fig. 5D (Excit-1 enhancer), mice were randomly assigned to receive either CNO or vehicle prior to CQ injection; these same mice were subsequently assigned to different chronic pain models (CFA or SNI) for separate experiments.

### Immunofluorescence (IF) and Fluorescence *In Situ* Hybridization (FISH)

#### Tissue Preparation

##### Mice

Mice were deeply anesthetized via intraperitoneal (i.p.) injection of ketamine (100 mg/kg) and xylazine (20 mg/kg) and transcardially perfused with ice-cold, RNase-free phosphate-buffered saline (PBS), followed by 4% paraformaldehyde (PFA) in RNase-free PBS. The lumbar spinal cord and corresponding dorsal root ganglia (DRGs) were harvested, post-fixed in 4% PFA overnight at 4°C, and then cryoprotected by sequential incubation in 20% and 30% sucrose solutions. Tissues were embedded in Optimal Cutting Temperature (OCT) compound, and transverse sections were cut on a cryostat (Leica CM1950) at 16 µm (for DRGs) or 20 µm (for spinal cord). Sections were mounted directly onto Superfrost Plus slides (Fisherbrand, 22-037-246) and stored at -80°C until use.

#### Macaques

3-year-old *Macaca mulatta* (M94-17) and 5-year-old *Macaca fascicularis* (M263-23) were deeply anesthetized with ketamine, followed by administration of sodium pentobarbital. M94-17 was transcardially perfused with ice-cold artificial cerebrospinal fluid (aCSF). Following perfusion, the lumbar segment of the spinal cord was harvested. Lumbar spinal cord was snap frozen and transverse sections were cut using a cryostat (Leica CM1950) at 20 µm as described above. M263-23 was transcrardially perfused with ice-cold RNAse-free PBS followed by 4% PFA in RNAse-free PBS. The lumbar spinal cord was harvested, post-fixed in 4% PFA overnight at 4°C, and then cryoprotected by sequential incubation in 20% and 30% sucrose solutions. Transverse spinal cord sections were prepared using a cryostat (Leica CM1950) as described above.

To reduce lipofuscin-induced autofluorescence in macaque spinal cord sections, a photobleaching protocol was employed as previously described (*81*). Briefly, slides containing tissue sections were immersed in ice-cold, RNase-free PBS in a plexiglass dish. The dish was placed on ice approximately 10 cm below a LED light source (Husky, HD12000DIM). The sections were then continuously illuminated for 48 hours at 4°C to photobleach endogenous fluorophores before FISH.

#### Immunofluorescence (IF) staining

Sections were brought to room temperature and washed in PBS. DRG sections were blocked for 1 hour at room temperature in PBS containing 10% normal donkey serum (NDS; Jackson ImmunoResearch Labs, 017-000-121) and 0.5% Triton X-100. Sections were then incubated overnight at 4°C with primary antibodies (see table S1 for details) diluted in PBS containing 5% NDS and 0.3% Triton X-100. The following day, sections were washed three times in PBS with 0.05% Tween-20 (PBST) and incubated for 2 hours at room temperature with secondary antibodies (table S1) diluted in PBS with 2% NDS and 0.3% Triton X-100. For spinal cord sections, the same procedure was followed, but a higher concentration of Triton X-100 (1%) was used in all blocking and antibody solutions to enhance penetration. Finally, all sections were washed three times in PBST and coverslipped using Fluoromount-G, with or without DAPI (Southern Biotech, 0100-20).

#### Fluorescence *in situ* hybridization (FISH)

FISH was performed on prepared spinal cord sections using the RNAScope Multiplex Fluorescent Reagent Kit v2 (ACD Bio, 323110) according to the manufacturer’s protocol for fixed-frozen tissue, with minor modifications as previously described (*82*). Specifically, Protease IV (ACD Bio, 322336) was used for the protease treatment step. A detailed list of all RNAScope probes and fluorophore reagents is provided in table S1.

For combined FISH and IF, the IF protocol was performed immediately following the final amplification step of the RNAScope assay. Sections were blocked for 1 hour at room temperature in Tris-buffered saline (TBS) containing 1% bovine serum albumin (BSA; TBS-BSA). Primary antibody incubation was performed overnight at 4°C in the same buffer, followed by three washes in TBS with 0.05% Tween-20 (TBST). Secondary antibody incubation was performed for 2 hours at room temperature in TBS-BSA. Finally, sections were washed three times in TBST and coverslipped using ProLong Gold Antifade Mounting medium with DAPI (Thermo Fisher Scientific, P36931).

#### Image acquisition and quantification

Images were acquired on a Nikon A1R confocal laser-scanning microscope equipped with 405, 488, 561, 640 nm excitation lasers and controlled by Nikon NIS-Elements software. All channels were acquired sequentially to prevent emission crosstalk. Z-stacks were captured at 20x magnification (with appropriate digital zoom to achieve Nyquist resolution) and stitched together. To reduce shot noise inherent to resonant scanning, the Denoise.ai function in NIS-Elements was applied to each channel. All quantitative analysis was performed on maximum intensity projections of these denoised z-stacks. For each animal, at least three spinal cord or DRG sections were imaged for quantification.

For quantification of viral expression, sections with the highest transgene expression and no signs of physical needle damage were selected for imaging. For cFos analysis, sections exhibiting the most robust cFos induction were chosen. Quantification was restricted to the relevant anatomical regions: the medial dorsal horn (laminae I-II) of lumbar segments L3-L5 for hindpaw-related stimuli, and the middle dorsal horn (laminae I-II) of segment L3 for calf-related stimuli.

Quantification of FISH and IF signals was performed using the Cell Counter plugin in Fiji/ImageJ. An experimenter blinded to the experimental conditions performed cell counts whenever possible. For FISH analysis, cells were considered positive if the number of puncta co-localized with a DAPI-stained nucleus exceeded a defined threshold above background (≥10 puncta for *cFos*; ≥3 puncta for *Tac1*, *Slc32a1*, *Pdyn*, *Pvalb, SKOR2,* and *LMO3*; and ≥2 puncta for *Grpr*. Data quantified from least three sections were averaged for number of cells (e.g. number of hM4Di^+^ cells) or were summed and normalized to percent of hM4Di (e.g. *Grpr^+^Tac1^+^* cells with hM4Di/total hM4di^+^ cells) or percent of cell type expressing hM4Di (e.g. *Grpr^+^Tac1^+^* cells with hM4Di/total *Grpr^+^Tac1^+^* cells).

#### Treadmill induced *cFos* analysis

To assess neuronal activation following innocuous mechanical stimulation of the hindpaws, mice were subjected to treadmill walking as previously described (*31*) with minor modifications. Mice were first habituated to the testing room for 30 minutes, followed by a 5-minute acclimation period within individual lanes of a multi-lane rodent treadmill (Columbus Instruments, Exer-6M). The treadmill speed was then gradually increased from stationary to 10 cm s^-1^ over a 5-minute period. Mice walked at this final speed for 15 minutes and were subsequently returned to their home cages. Care was taken to minimize any further hindpaw stimulation. For experiments involving chemogenetic manipulation, CNO (5mg kg^-1^, i.p.) or vehicle (saline, i.p.) was administered 30 minutes before the start of the 15-minue walking period at the final speed. Mice were perfused, between 30-60 minutes after the completion of treadmill walking, and spinal cords were collected for FISH analysis as described above.

#### Chloroquine induced *cFos* analysis

To examine neuronal activation following a pruritogenic stimulus, mice were briefly anesthetized with 2.5% inhaled isoflurane and received an i.d. injection of CQ (5 μL of 20 mg mL^-1^ in sterile saline) or vehicle (5 μL of sterile saline) into the calf, as described for behavioural assessments. Following injection, mice were maintained under light anesthesia (1-1.5% inhaled isoflurane) for 30 minutes to prevent biting-induced *cFos* expression. After this period, anesthesia was discontinued, and mice were returned to their home cages. No calf-directed biting was observed upon recovery. In Fig. 3, J to L, all mice received CNO (5 mg kg^-1^, i.p.) 30 minutes before the i.d injection of CQ. Spinal cord samples were collected for FISH analysis 45-60 minutes after the CQ or vehicle injection, as described above.

#### Analysis of single nucleus RNA sequencing (snRNA-seq) database

For snRNA-seq analysis, we utilized the publicly available harmonized mouse spinal cord atlas dataset from Russ et al. (2021) (*25*), which includes data from both juvenile and adult mice. Data processing and analysis were performed using the Seurat R package (5.1.0). Raw count data from both juvenile and adult datasets were normalized using the NormalizeData() function. The original cluster annotations were used for the subsequent analysis. Nuclei with > 0 expression level for *Grpr* were selected, and the distribution of these *Grpr^+^*nuclei across the annotated excitatory neuron clusters was visualized (fig. S4A). Differentially expressed genes (DEGs) between the Excit-14 and Excit-16 clusters, specifically within the *Grpr^+^* nuclei, were identified using the FindMarkers() function. Results were visualized as an EnhancedVolcano plot (fig. S2B).

### Spatial transcriptomics

#### Xenium assay

Xenium spatial transcriptomics experiments were performed as previously described (*26*). Mice were deeply anesthetized with 100 mg kg^-1^ ketamine and 20mg kg^-1^ xylazine i.p. and transcradially perfused with ice cold RNAse-free PBS. Lumbar spinal cord segment (L3-L5) was dissected and embedded in optimal cutting temperature embedding medium (O.C.T.; Fisherbrand, 23-730-571) and frozen following the <u>Xenium In Situ for Fresh Frozen Tissues -</u> <u>Tissue Preparation Guide</u>. Lumbar spinal cords were sectioned at 10 μm, with at least 40 μm between each section, and mounted onto Xenium slides with Leica CM1950 cryostat. Slides were stored -80 ℃ until the assay was performed. The spatial transcriptomics assay was performed with Mouse brain panel with 50 custom add-on genes (table S2) on the Xenium analyzer following the manufacturer’s protocol.

#### Xenium cell labeling and quantification

Neurons were label based on a previously described and validated approach (*26*) with minor modifications. A mouse snRNA-seq dataset from Russ et al. (2021) (*25*), previously ascribed species-conserved neuron subtype labels from Arokiaraj et al. (2024) (*26*), was used as a reference to annotate the Xenium dataset. The reference was hierarchically organized into three annotation levels: major cell type, neuron subclass, and neuron subtype. At the cell type level, cells were broadly categorized as neuron or various glial and immune cell types. Within the neuronal subclass, neurons were divided into mid/ventral versus dorsal horn versus Exc-PDYN. Neuron subtypes included mid/ventral excitatory, inhibitory, and motor neurons, and the dorsal subtypes as described previously (*26*). Xenium cells were labeled based on iterative integrations across the three annotation levels with the reference snRNA-seq described above. snRNA-seq genes were limited to those also measured by Xenium, and both assays were log-normalized. First, labels were transferred at the major cell type level using Seurat v5’s FindIntegrationAnchors() and IntegrateData() based on canonical correlation analysis (CCA), and finally TransferLabels(). CCA was used because of major differences in the dynamic range of measured genes for snRNA-seq versus Xenium. Following cell-type-level annotation, both the reference and query datasets were subsetted to neurons. A second round of integration was conducted to assign neuron subclass annotations. This process was repeated once more within each subclass subset to assign precise neuron subtypes. Finally, only cells with an integration predicted confidence score above 0.5 were kept for further analysis. Cells with > 0 *Grpr* transcripts and cells with > 3 *Tac1* transcripts were considered positive. Representative figures were generated by Xenium Explorer (Version 3.0.0, 10x Genomics).

#### Targeted ablation of spinal *Grpr^+^* neurons

To preferentially ablate bombesin-responsive *Grpr^+^* neuron, we used a two-step viral and neurotoxin approach. First, homozygous Grpr^Cre^ mice were intraspinally injected with AAV8.hSyn.DIO.hM4Di-mCherry to express inhibitory DREADDs in all *Grpr*^+^ neurons, as described above. Two weeks after AAV injection, mice received a single 6 μL intrathecal (i.t.) injection of either bombesin-saporin (300ng; Advanced Targeting Systems, IT-40) or blank-saporin (300ng; Advanced Targeting Systems, IT-21) as a control. The i.t. injections were performed between L5 and L6 vertebrae in lightly anesthetized mice using a 30-gauge needle attached to a Hamilton syringe. Chemical itch assay was performed 2 weeks following the i.t injections (Fig. 4G). The same mice that received CQ in Fig. 4g were subjected to CFA model 7 days after the chemical itch experiments (Fig. 4H). For the long-course STZ and HFD models, bombesin- or blank-saporin was injected 2 weeks prior to behavioural analysis to prevent ablation associated plasticity in the spinal cord. As for the HFD model, the same mice from Fig. 1i were randomly divided into bombesin- or blank-saporin group.

#### Targeted recombination in active populations (TRAP2)

To selectively capture and manipulate bombesin-responsive neurons, we utilized homozygous TRAP2 mice, which express a tamoxifen-inducible CreERT2 recombinase from the *cFos* promoter (*56*). TRAP2 mice were first intraspinally injected with AAV8.hSyn.DIO.hM4Di-mCherry. Three weeks following the intraspinal AAV injection, mice received a single i.p. injection of 4-hydroxyamoxifen (4-OHT; 50 mg kg^-1^; Sigma-Aldrich, H6278) dissolved in Kolliphor EL (Sigma-Aldrich, C5135). Immediately following the 4-OHT injection, neuronal activity was induced by an i.t. injection of bombesin (0.1 nmol; Anaspec, AS-20665), dissolved in sterile PBS, to activate *Grpr*-expressing neurons. In parallel, a control group received an i.t. injection of sterile PBS. Mice were maintained under light anesthesia (1-1.5% inhaled isoflurane) for 60 minutes to prevent bombesin induced scratching behaviour. Mice were returned to home cage after recovery from isoflurane. Behavioural and histological experiments were performed at least two weeks after to allow for robust hM4Di-mCherry expression in the TRAPed population. Chemical itch was first assessed with both PBS and bombesin TRAPed mice receiving CNO (5 mg kg^-1^, i.p.). Then, PBS or bombesin TRAPed mice were randomly assigned to CFA or long-course STZ groups. A group of bombesin or PBS TRAPed mice, which chemical itch was not assessed, was also subjected to CFA or long-course STZ models. Mechanical thresholds were measured as described previously.

#### Statistical analysis

All statistical analysis were performed using GraphPad Prism (version 10, GraphPad Softwrae). Data are reported as mean ± standard error of the mean (s.e.m.) unless otherwise stated in the figure legends. The specific statistical tests and number of animals (n) are detailed in the corresponding figure legends. A P-value less than 0.05 was considered statistically significant. Significance is denoted as *P < 0.05, **P < 0.01, ***P < 0.001, ****P < 0.0001. Sample sizes for mouse behaviour experiments were determined by power analysis with significance level of 0.05, and a power of 80% using the data previously generated by our laboratory (*31, 32*). No explicit sample size determination was made for FISH and IF experiments; however, our previous work (*31, 32*) was used to estimate sample sizes. Mechanical sensitivity data were analyzed over time using repeated-measures one-way or two-way ANOVA, followed by Bonferroni’s post-hoc test for multiple comparisons between treatment groups at specific time points. For single time-point comparisons between two groups, unpaired two-tailed Welch’s t-test was used. For paired comparisons within the same animals before and after treatment, a paired two-tailed t-test was utilized. Comparisons of cell counts or percentages between two independent groups were performed with unpaired two-tailed Welch’s t-test. For comparisons involving three or more groups (e.g. distribution of hM4Di^+^ cells across *Grpr^+^* neuron subtypes), two-way ANOVA followed by Bonferroni’s post-hoc test was used. All n values for histological data represent the number of animals. DEGs between clusters was identified using the non-parametric Wilcoxon Rank Sum test, as implemented in Seurat FindMarkers function. P-values were adjusted for multiple comparisons using the Bonferroni correction. Detailed summary of statistical analysis is provided in table S3.

## Acknowledgments

We thank August Henry, Kailey Go, Kelsey Go, Varsha Sriram, Rachelle Huynh, Bennett Allisson, Zane Hamdan, Nikolka Majernikova, An Binh Tran, and Amber Frantz for technical assistance, behaviour scoring, and cell counting. We thank Drs. William Stauffer and David Lewis for providing the macaques used in this study. Drs. David Ginty and Victoria Abraira provided the PKCγ^CreERT2^ mice and Dr. Carmen Birchmeir provided the Lbx1^Cre^ mice. AAV8.nEF.Con/Fon.hM3Dq-mCherry was kindly gifted to us by Dr. Karl Deisseroth.

## Funding

National Institutes of Health grant R56NS133364 (ARP and RPS)

National Institutes of Health grant R56NS104964 (RPS)

National Institutes of Health grant R01NS107364 (RPS)

National Institutes of Health grant R01NS139529 (RPS)

## Author contributions

M-cN and RPS conceived the study. M-cN, CP, and XZ performed intraspinal injections. M-cN, KAC, S-PGW, C.P, and DJH generated the pain models. KAC and S-PGW performed the mechanical allodynia assessments. M-cN and KAC performed the Xenium spatial transcriptomics. M-cN, KAC, and DJH performed the chemical itch assays. M-cN performed intrathecal injections. M-cN, KAC, DJH, and MG performed the IF and FISH. M-cN and RPD collected macaque spinal cords. M-cN and KAC executed TRAP2 experiments. M-cN, KAC, and DJH performed the treadmill and chloroquine *cFos* experiments. M-cN and KAC executed nocifensive behaviour assay. M-cN acquired the confocal images. M-cN, RPS, ARB, and RvdW designed the viral constructs. MJL, BNP, DY, and ARP developed and executed machine learning pipeline for enhancer identification and prioritization. MJL, RvdW, ARB, BNP, and ARP provided Excit-1 and SKOR2.103 sequences. M-cN, KAC, S-PGW, CP, MJL, SB, DJH, MG, SL, HM, and RPS analyzed the data. M-cN and RPS wrote the manuscript.

## Competing interests

The authors declare the following competing interests: ARP is founder and CEO of Snail Biosciences. ARP and BNP have Snail Biosciences stockholdings. The authors declare that the research was conducted in the absence of any commercial or financial relationships that could be construed as a potential conflict of interest.

## Data and materials availability

The authors declare that the data supporting the findings of this study are available within the paper and its supplementary information files.

---

**Fig. S1.**
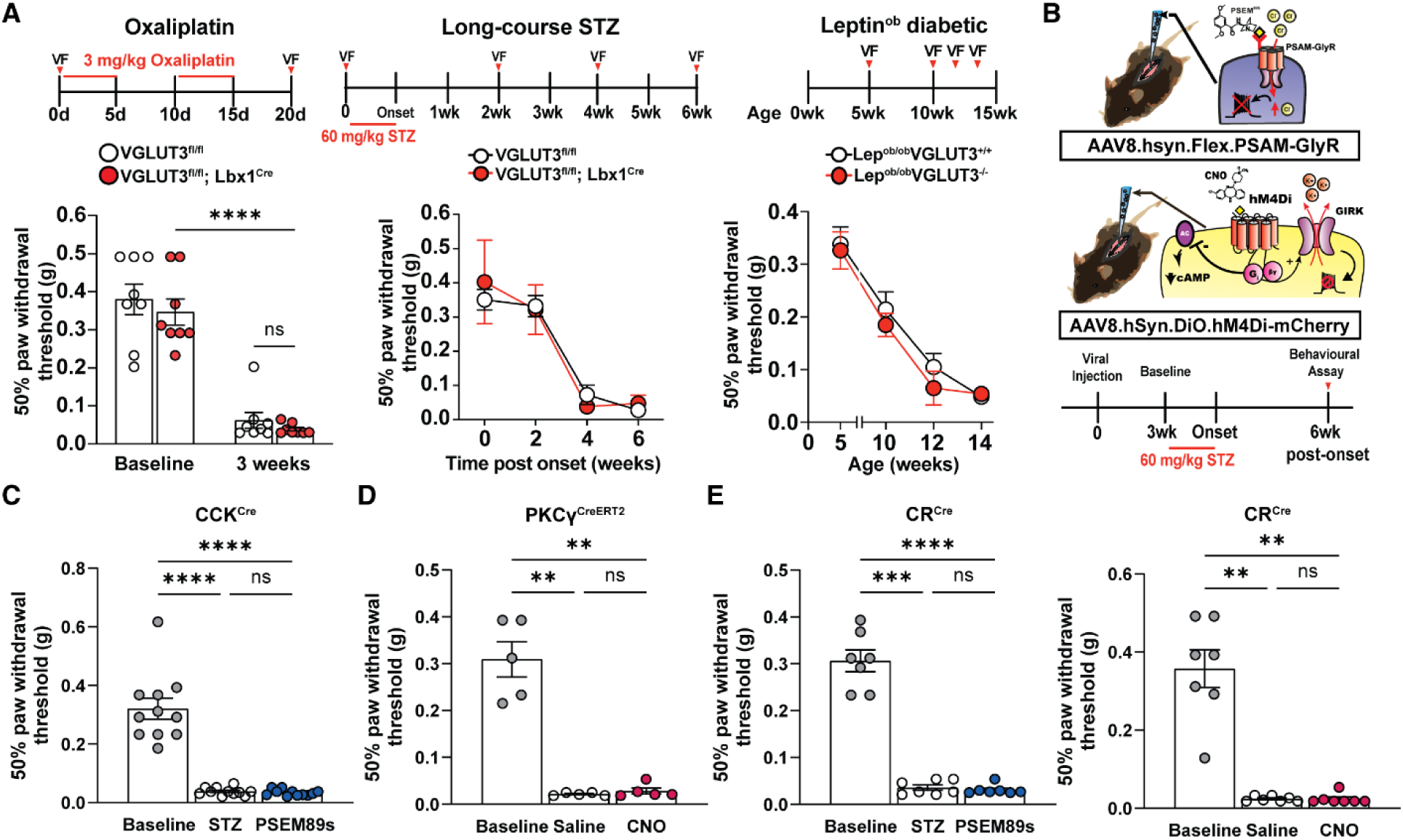
Mechanical allodynia circuits for polyneuropathic injuries are distinct from inflammatory and neuropathic injuries. **(A)** 50% paw withdrawal threshold in control and VGLUT3 knockout mice at baseline and after polyneuropathic injury. Oxaliplatin model (n=8 per group), long-course streptozotocin (STZ) (control, n=3; knockout, n=4), and leptin deficiency (ob/ob) model of diabetes (control, n=4; knockout, n=9). **(B)** Schematic of chemogenetic inhibition strategy to assess the role of specific dorsal horn neuron populations in the long-course STZ model of diabetic neuropathy. (**C to E),** 50 % paw withdrawal threshold at baseline, after long-course STZ, and after chemogenetic inhibition of **(C)** CCK neurons (n=11), (D) PKCγ neurons (n=5), and (E) calretinin (CR) neurons (n=7). No statistically significant changes to von Frey thresholds were observed following chemogenetic inhibition of CCK, PKCγ, and CR neurons. All data are mean ± s.e.m. Statistical analysis was performed using two-way ANOVA with Bonferroni’s post hoc test (oxaliplatin), repeated-measures two-way ANOVA with Bonferroni’s post hoc test (long-term STZ and leptin deficiency), and repeated-measures one-way ANOVA with Bonferroni’s post hoc test (C to E). **P < 0.01, ***P < 0.001, ****P < 0.0001.

**Fig. S2.**
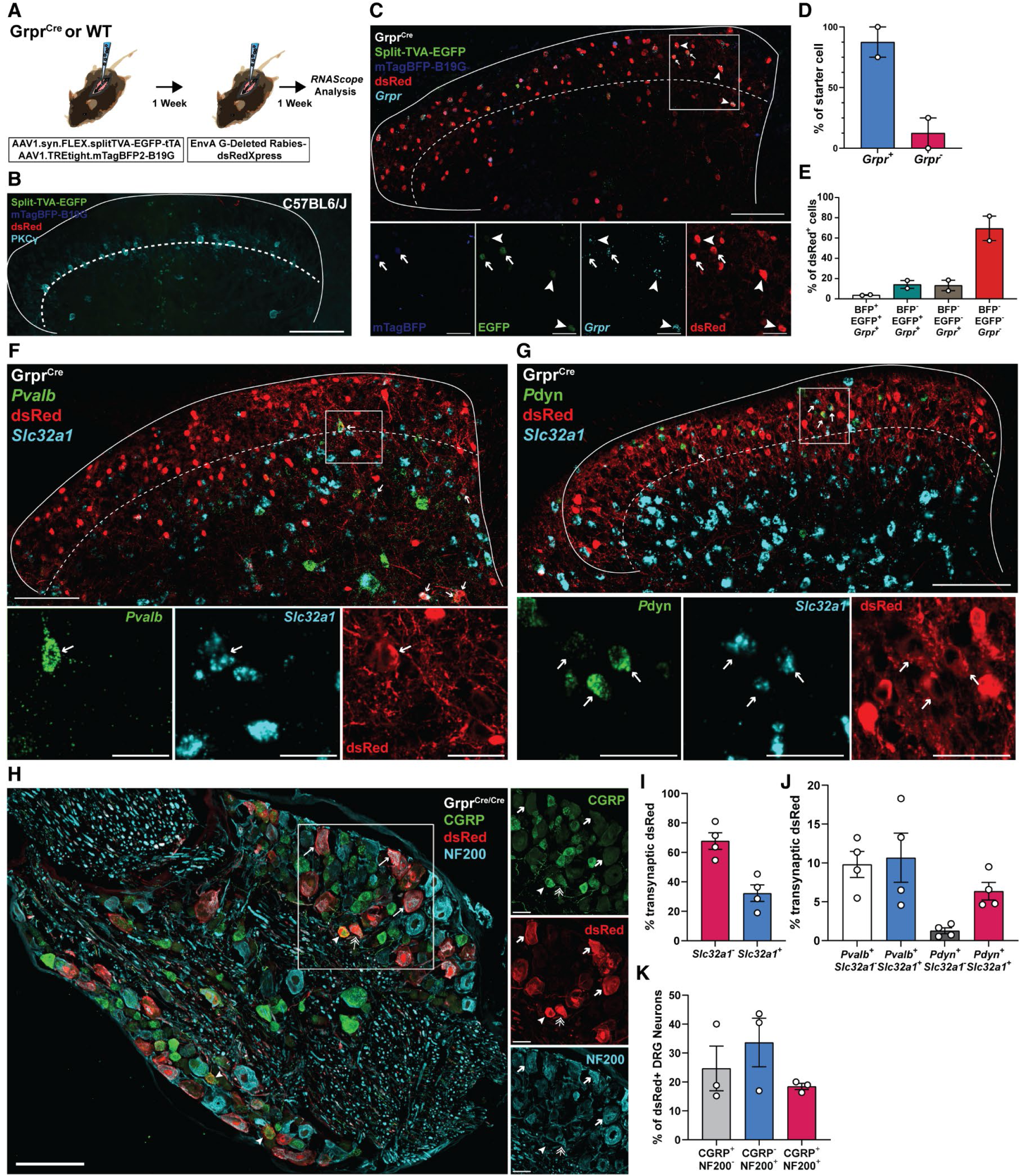
Monosynaptic rabies tracing reveals synaptic inputs to *Grpr^+^* neurons. **(A)** Experimental schematic for monosynaptic rabies tracing in homozygous Grpr^Cre^ or wild type (WT) mice. **(B)** Lumbar dorsal horn section from a WT mouse showing TVA (green), B19G (blue), dsRed (red), and PKCγ (cyan) expression after intraspinal delivery of helper AAVs and EnVA G-deleted rabies. **(C)** Lumbar dorsal horn section from a Grpr^Cre^ mouse showing TVA (green), B19G (blue), dsRed (red), and *Grpr* (cyan) expression after intraspinal delivery of helper AAVs and EnVA G-deleted rabies. Arrowheads indicate cells with TVA, dsRed and *Grpr*. Arrows indicate cells with TVA, B19G, dsRed, and *Grpr* (starter cells). **(D)** Percentage of starter cells (TVA^+^, B19G^+^, and dsRed^+^) with or without *Grpr* transcript (n=2 mice). **(E)** Percentage of dsRed^+^ cells with or without TVA, B19G, and *Grpr* (n=2 mice). **(F and G)** Lumbar dorsal horn section showing dsRed (red), *Slc32a1* (cyan), and (F) *Pvalb* or (G) *Pdyn* (green) expression following monosynaptic rabies tracing in Grpr^Cre^ mice. Arrows indicate dsRed cells with *Slc32a1* and (F) *Pvalb* or (G) *Pdyn* expression. **(H)** Lumbar dorsal root ganglia (DRG) section showing CGRP (green), dsRed (red), and NF200 (cyan) expression following monosynaptic rabies tracing in Grpr^Cre^ mice. Arrows indicate dsRed cells with NF200, arrowheads indicate dsRed cells with CGRP, and double arrows indicate dsRed cells with both NF200 and CGRP. **(I to K)** Percentage of transsynaptic dsRed^+^ cells (TVA^-^) following monosynaptic rabies tracing in Grpr^Cre^ mice in (I) excitatory (*Slc32a1^-^*) and inhibitory (*Slc32a1^+^*) cells in the dorsal horn, (J) inhibitory and excitatory *Pvalb^+^* and *Pdyn^+^*cells in the dorsal horn, and CGRP^+/-^ and NF200^+/-^ DRG neurons (n=4, 4, and 3 mice). Data are mean ± s.e.m. Dashed lines in the representative figures **(B, C, F, and G)** delineate the lamina II/III borders. Scale bars: 100 μm (main images), 30 μm (insets).

**Fig. S3.**
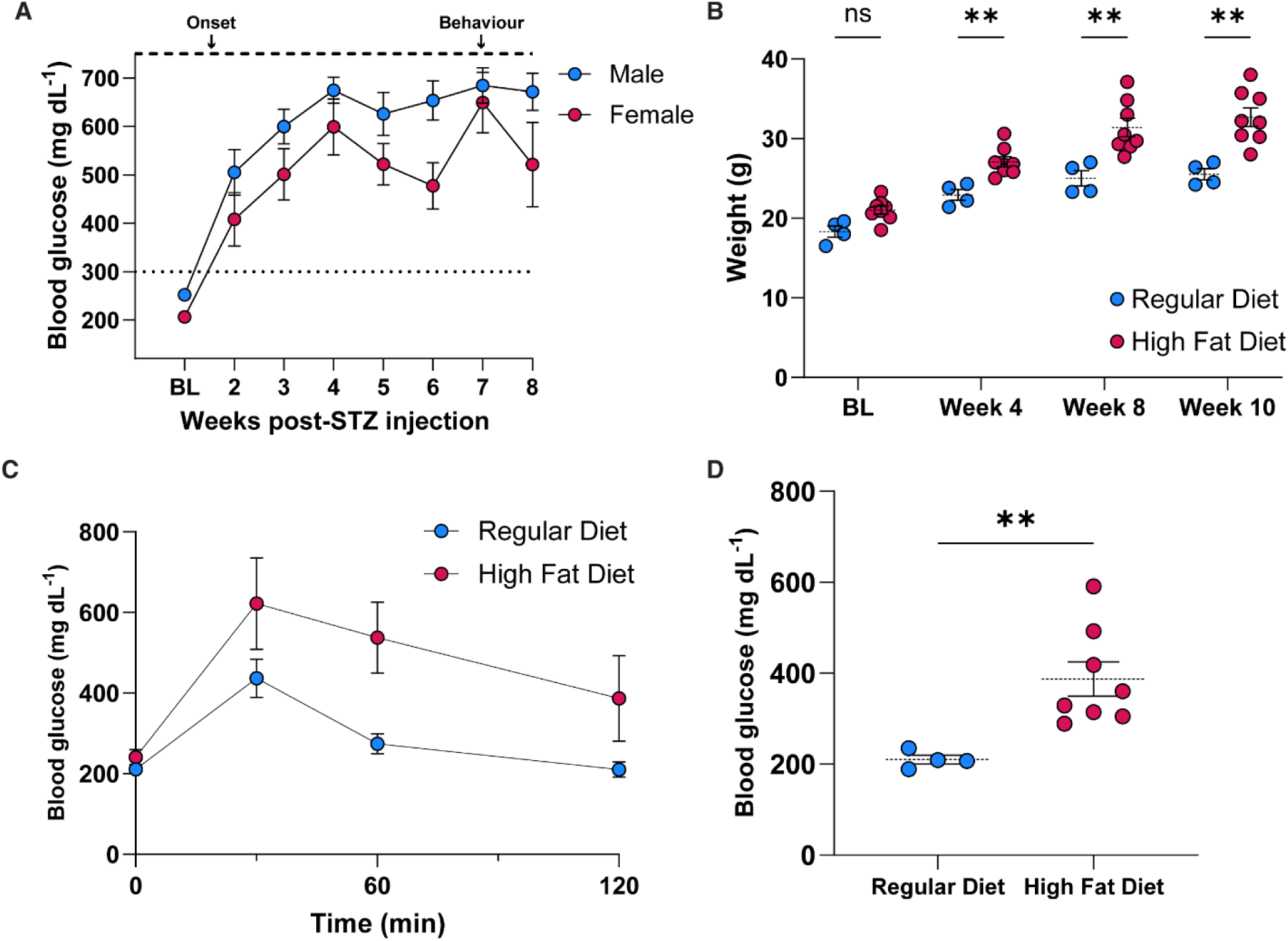
Long-course STZ and HFD model causes hyperglycemia and glucose intolerance respectively. **(A)** Blood glucose levels following intraperitoneal injection of STZ across experimental time course (n=7 males, and n=5 females). Dotted lines show threshold for hyperglycemia and dashed lines represent maximum measurable blood glucose levels. **(B)** Weights of regular or high fat diet fed male mice across 10 weeks (n=4 regular diet, n=8 high fat diet). **(C)** Blood glucose levels in male mice fed regular or high fat diet after 10 weeks. Data are mean ± standard deviation of the mean (s.d.) **(D)** Blood glucose levels following a systemic glucose injection in mice (C) after 120 minutes (n=4 regular diet, n=8 high fat diet). Data are mean ± s.e.m. unless stated otherwise. Statistical analysis was performed using repeated-measures two-way ANOVA with Bonferroni’s post-hoc test (B) and unpaired two-tailed Welch’s t-test (D). **P < 0.01

**Fig. S4.**
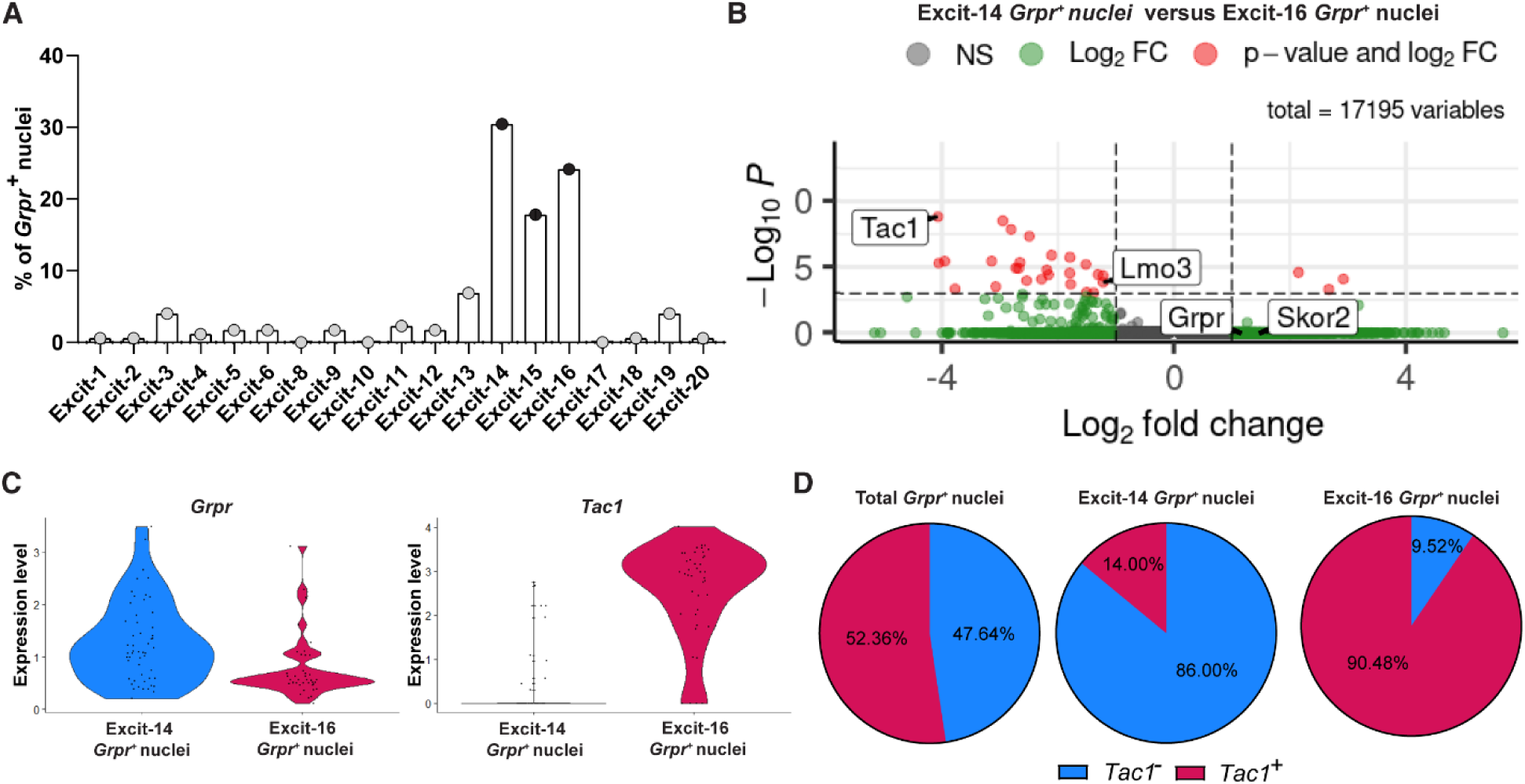
*Grpr^+^* neurons segregate into transcriptionally distinct clusters distinguished by *Tac1* expression. **(A)** Distribution of *Grpr^+^* nuclei across excitatory neuron clusters in the harmonized mouse spinal cord atlas of Russ *et al.* (2021). **(B)** Volcano plot showing differential gene expression between *Grpr^+^*nuclei in the Excit-14 and Excit-16 clusters. Significantly enriched genes (red circles) were defined as those with -log10 P > 3 and log2 fold change >1. Negative log2 fold change values indicate enrichment in the Excite-16 cluster; positive values indicate enrichment in the Excit-14 cluster. **(C)** Violin plots illustrating the relative expression levels of *Grpr* (left) and *Tac1* (right) transcripts within *Grpr^+^* nuclei belonging to the Excit-14 and Excit-16 clusters. **(D)** Proportion of *Grpr^+^* nuclei expressing *Tac1* transcripts, shown for all *Grpr^+^* nuclei (left), *Grpr^+^* nuclei within the Excite-14 cluster (middle), and *Grpr^+^* nuclei within the Excite-16 cluster (right).

**Fig. S5.**
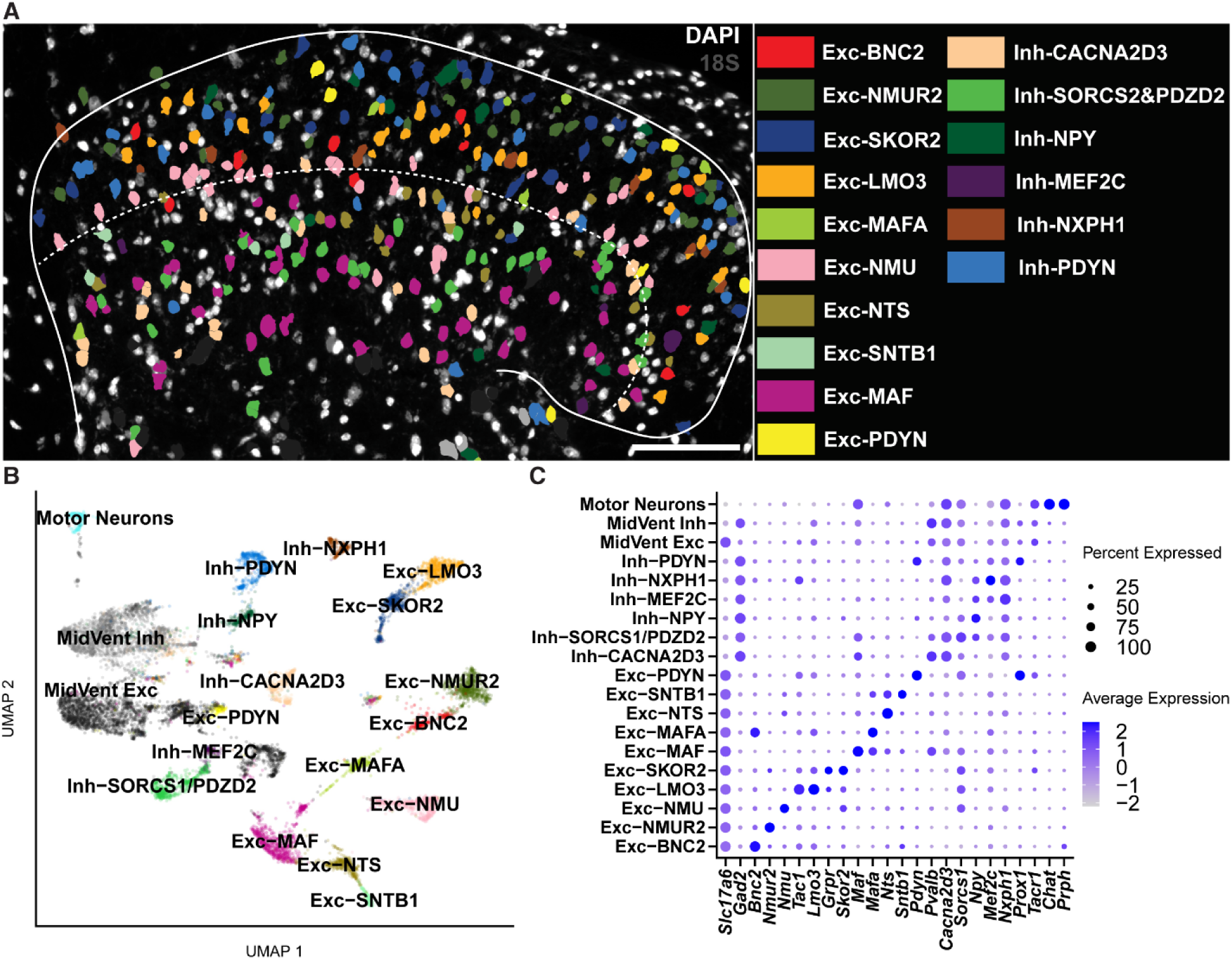
Species-conserved dorsal horn neuron populations detected by Xenium spatial transcriptomics. **(A)** Xenium spatial transcriptomics image of lumbar dorsal horn section showing species-conserved dorsal horn neuron subtypes. Same image as in main Fig. 2B with all dorsal horn neuron subtypes. Dashed lines delineate lamina II/III border. Scale bar: 100 μm. **(B)** UMAP showing species conserved dorsal horn neuron subtypes detected by Xenium spatial transcriptomics platform (n=3 mice). **(C)** Dot plot of marker gene expression patterns of dorsal horn neuron subtypes detected by Xenium spatial transcriptomics platform (n=3 mice).

**Fig. S6.**
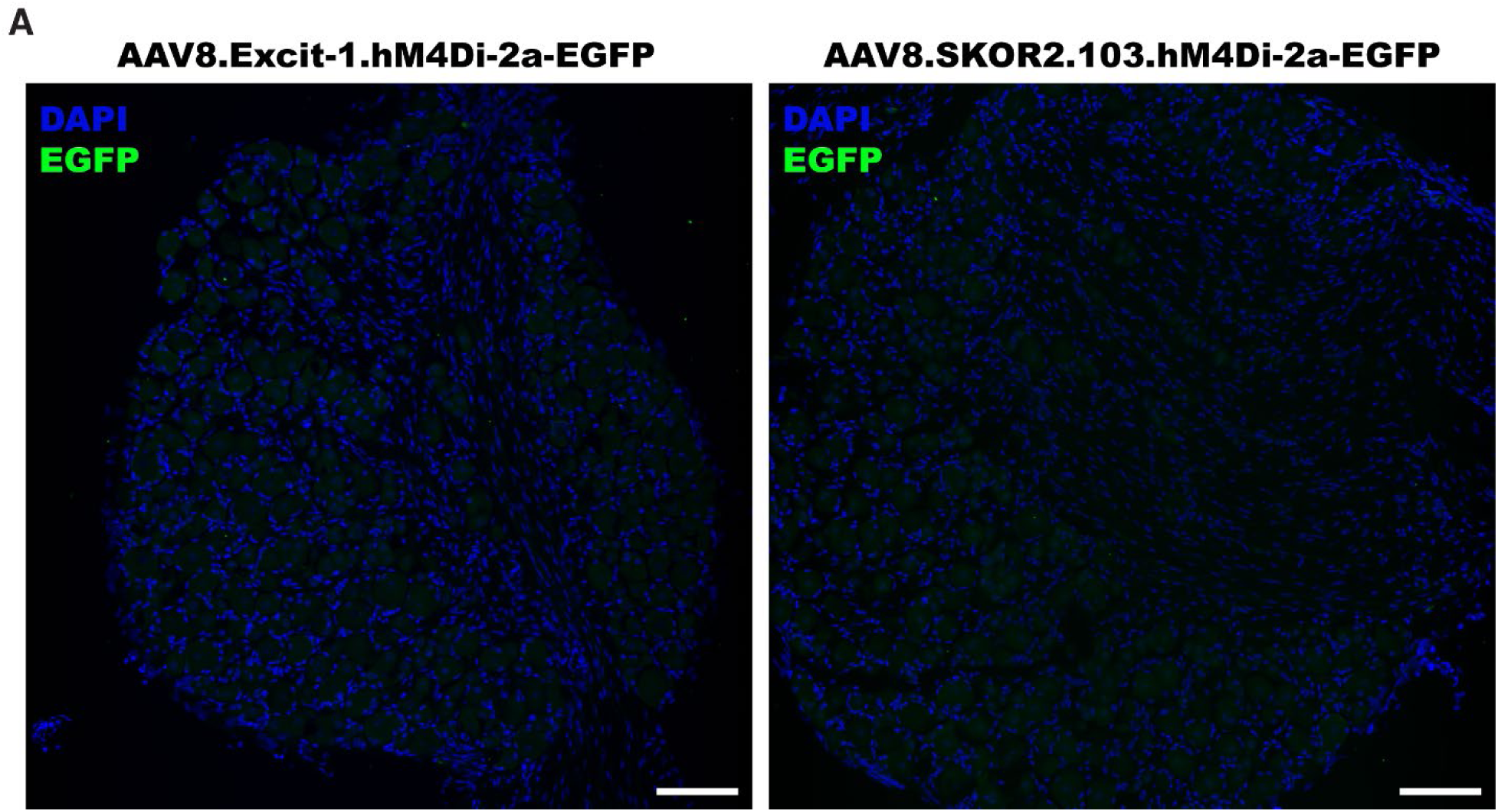
Excit-1 and SKOR2.103mBG enhancer elements do not transduce dorsal root ganglia neurons. **(A)** Confocal images of L4 DRG sections from mice intraspinally injected with AAV8.Excit-1.hM4Di-2a-EGFP (left) or AAV8.SKOR2.103.hM4Di-2a-EGFP (right). Immunofluorescence staining for GFP (Green) showed no detectable fluorescence in L3-L5 DRGs (n=3 mice each). Scale bar: 100μm.

**Fig. S7.**
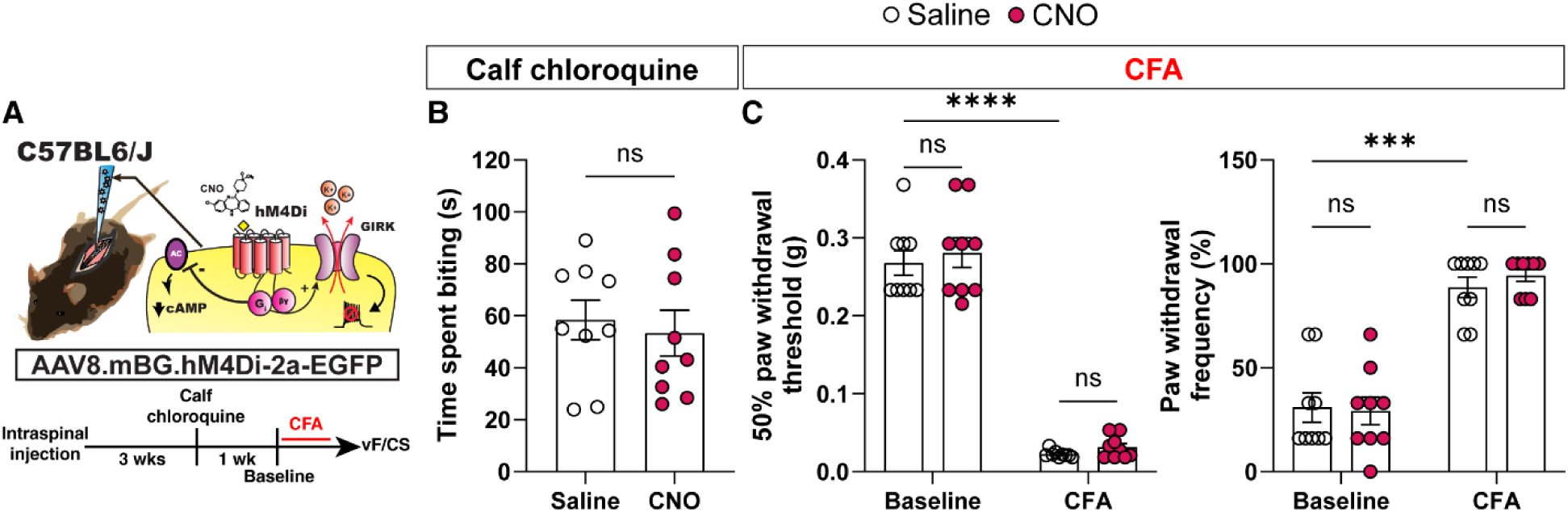
Chemogenetic inhibition of beta-globin minimal promoter-targeted neurons does not affect chemical itch or mechanical allodynia. **(A)** Experimental strategy to test the role of mBG promoter-targeted neurons in chemical itch and mechanical allodynia. **(B)** Itch-like behaviour after intradermal chloroquine, with saline or CNO in WT mice intraspinally injected with AAV8.mBG.hM4Di-2-EGFP (n=9 mice). (**C)** 50% paw withdrawal threshold (von Frey test, left) and frequency (cotton swab test, right) with saline or CNO in mice before and after CFA (n=9 mice; same mice as (B)). Data are mean ± s.e.m. Statistical analysis was performed with paired two-tailed t-test (B), and repeated-measures two-way ANOVA with Bonferroni’s post-hoc test (C). ***P < 0.001, ****P < 0.0001.

## Notes

### Summary of Updates

Figure resolution was poor. Re-uploaded figures to improve quality.

